# TWO DISULFIDE-REDUCING PATHWAYS ARE REQUIRED FOR THE MATURATION OF PLASTID *C*-TYPE CYTOCHROMES IN *CHLAMYDOMONAS REINHARDTII*

**DOI:** 10.1101/2022.10.14.512171

**Authors:** Ankita Das, Nitya Subrahmanian, Stéphane T. Gabilly, Ekaterina P. Andrianova, Igor B. Zhulin, Ken Motohashi, Patrice Paul Hamel

**Author notes:** **Correspondence** Patrice P. Hamel, The Ohio State University, Department of Molecular Genetics and Department of Biological Chemistry and Pharmacology. 500 Aronoff Laboratory, 318 W. 12^th^ Avenue, Columbus, OH, 43210, USA. University of Florida, Department of Neurology, 1275 Center drive, Gainesville, Florida, USA. Calixar, Bâtiment Laennec, 60 avenue Rockfeller, Lyon, France (Stéphane T. Gabilly).

## Abstract

In plastids, conversion of light energy into ATP relies on cytochrome *f*, a key electron carrier with a heme covalently attached to a C*XX*CH motif. Covalent heme attachment requires reduction of the disulfide bonded C*XX*CH motif by CCS5 and CCS4, a protein of unknown function. CCS5 receives electrons from the oxido-reductase CCDA at the thylakoid membrane. In *Chlamydomonas reinhardtii*, loss of CCS4 or CCS5 function yields a partial cytochrome *f* assembly defect. Here we report that the Δ*ccs4ccs5* double mutant displays a synthetic photosynthetic defect due to a complete loss of holocytochrome *f* assembly, a phenotype that can be chemically corrected by reducing agents. In Δ*ccs4*, the CCDA protein accumulation is decreased, indicating that one function of CCS4 is to stabilize CCDA. Dominant suppressor mutations mapping to the *CCS4* gene were identified in photosynthetic revertants of the Δ*ccs4ccs5* mutants. The suppressor mutations correspond to changes in the stroma-facing domain of CCS4 and restore holocytochrome *f* assembly above the residual levels detected in Δ*ccs5*. Because disulfide reduction via CCS5 no longer takes place in Δ*ccs5*, we hypothesize the suppressor mutations enhance the supply of reducing power independently of CCS5, uncovering the participation of CCS4 in a distinct redox pathway. CCS4-like proteins occur in the green lineage and are related to mitochondrial COX16, a protein involved in a disulfide reducing pathway. We discuss the operation of two pathways controlling the redox status of the heme-binding cysteines of apocytochrome *f* and the possible function of CCS4 as a shared component between the two pathways.

Graphical abstract.
The Δ*ccs4ccs5* mutant exhibits a photosynthetic growth defect due to a complete loss of cytochrome *c* assembly.
Reduction of apocytochrome *f* in the thylakoid lumen requires the provision of reducing power through two different pathways, pathway 1 and 2. CCDA and CCS5, components of pathway 1, deliver electrons from stroma to apocytochrome *f* via thiol – disulfide exchange. CCS4 is involved in pathway 1 by stabilizing CCDA, but also functions through a CCS5 – independent pathway (pathway 2). In the absence of CCS5, gain – of – function mutations in the C terminus of CCS4 (indicated by a yellow star) enhance the delivery of reducing power either via CCDA or independently of CCDA to yet-to-be-discovered reductases

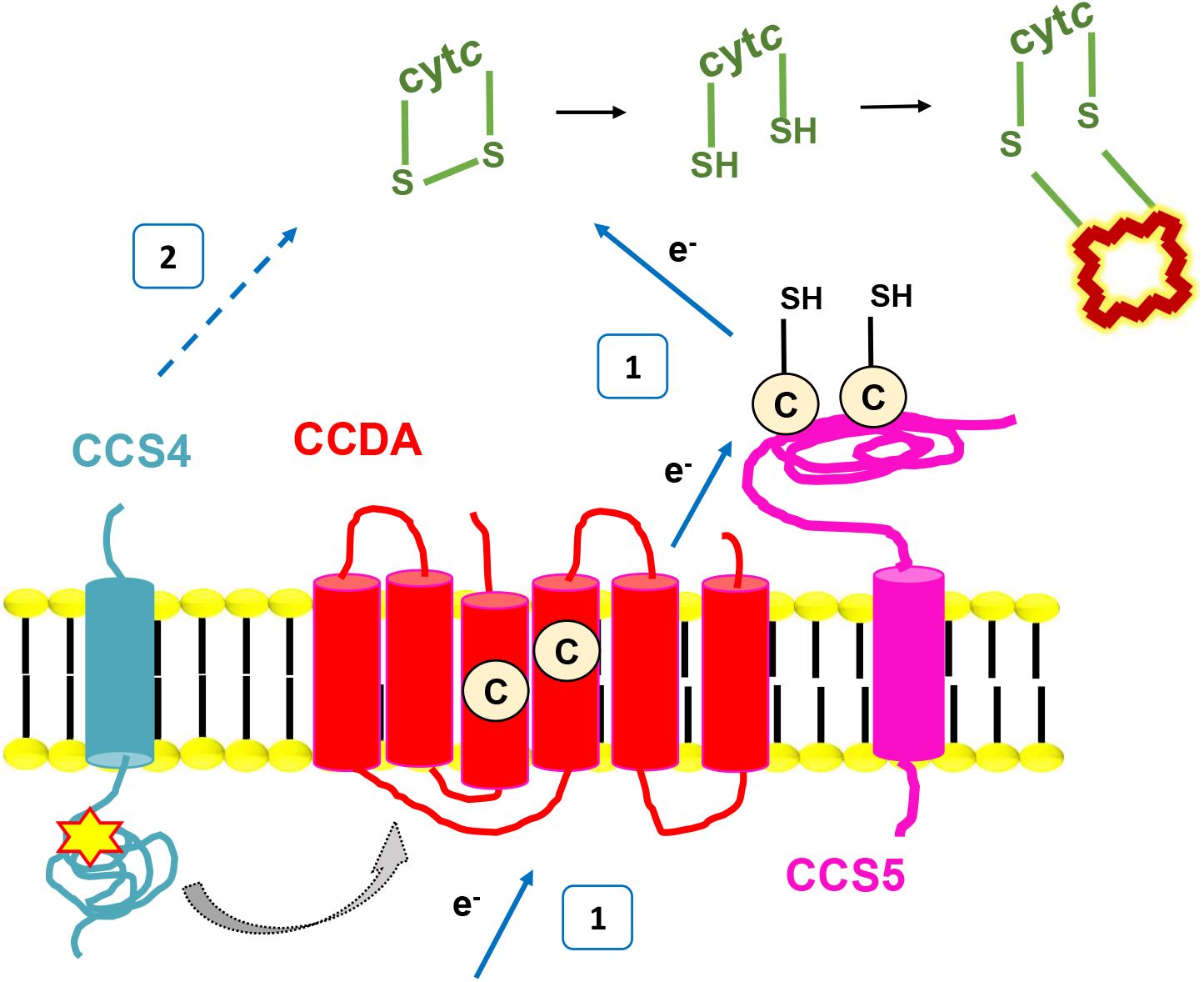

## Introduction

Cytochromes *c* or *c*-type cytochromes are ubiquitous heme (ferroprotoporphyrin IX) containing proteins that function as one-electron carriers in energy-transducing membranes in bacteria, archaea, mitochondria and plastids (Bertini *et al*. 2006; Ma and Ludwig 2019). Cytochromes *c* located on the positive side (or *p*-side) of energy-transducing membranes (*i*.*e*., the bacterial or archaeal periplasm, the thylakoid lumen in the plastid and mitochondrial intermembrane space) carry a heme moiety covalently bound to two (or rarely one) cysteines in a typical C*XX*CH motif called the heme-binding site on the apoprotein (Alvarez-Paggi *et al*. 2017). The histidine residue acts as an axial ligand to the iron in the heme moiety (Dickerson *et al*. 1971). Conversion of apocytochrome *c* to its corresponding holoform requires the formation of the thioether linkage(s) between the vinyl carbons in heme and the cysteine thiol(s) in the heme-binding motif. The chemical requirements for thioether bond formation were formulated from the *in vitro* reconstitution of holocytochrome *c* formation and the extensive genetic and biochemical dissection of cytochrome *c* maturation in bacterial and eukaryotic model systems (Cline *et al*. 2016; Das and Hamel 2021) In energy-transducing membranes, apo- to holocytochrome *c* conversion occurs on the *p*-side of the membrane and requires minimally: 1) the independent transport of apocytochrome *c* and heme substrates, 2) the chemical reduction of a) ferriheme to ferroheme and b) disulfide bonded heme - linking cysteines to free thiols, and 3) the stereospecific ligation of heme to the free thiols of the C*XX*CH motif via formation of thioether bond linkages. *In vivo*, heme ligation to C*XX*CH cysteine thiols is a catalyzed process requiring at least four different maturation systems (Systems I, II, III and IV) which are defined by signature assembly factors (Mavridou *et al*. 2013; Travaglini-Allocatelli 2013; Babbitt *et al*. 2015; Gabilly and Hamel 2017; Belbelazi *et al*. 2021). The diversity in cytochrome *c* assembly routes challenges our understanding of the process considering we view the stereospecific attachment of cysteine thiols to the vinyl carbons of heme as a simple chemical reaction (Bowman and Bren 2008).

In plastids and several bacteria, cytochrome *c* maturation is assisted by the CCS (CCS for <u>c</u>ytochrome *<u>c</u>* <u>s</u>ynthesis) factors, which were first identified through the isolation of the *ccs* mutants in the green alga *Chlamydomonas reinhardtii* (Gabilly and Hamel 2017; Das and Hamel 2021). The *ccs* mutants are specifically blocked in the attachment of heme to the apoforms of plastid cytochromes *c*, namely membrane-bound cytochrome *f* and soluble cytochrome *c*_6_. (Karamoko *et al*. 2013; Gabilly and Hamel 2017; Das and Hamel 2021). Cytochrome *f*, one of the plastid *c*-type cytochromes, is a structural subunit of the cytochrome *b*_6_*f* complex and an essential electron carrier in photosynthesis, (Kuras and Wollman 1994; Zhou *et al*. 1996). Consequently, the *ccs* mutants are deficient for cytochrome *b*_6_*f* assembly and photosynthesis (Cline *et al*. 2016; Gabilly and Hamel 2017).

In the thylakoid lumen, the redox state of the apocytochrome *f* C*XX*CH motif is under the control of a) CCS5, a thylakoid-bound disulfide reductase catalyzing the reduction of the disulfide bond formed between the cysteines of the heme-binding site into thiols and b) CCDA, a thylakoid membrane protein delivering reducing power from stroma to lumen (Lennartz *et al*. 2001; Page *et al*. 2004; Motohashi and Hisabori 2006; Gabilly *et al*. 2010; Motohashi and Hisabori 2010). CCS5 is classified as a member of the thioredoxin-like family, a diverse group of proteins including molecules able to mediate thiol-disulfide exchange reactions via a redox-active WC*XX*C motif (Atkinson and Babbitt 2009). CCS5 displays high similarity to membrane-anchored lumen-facing HCF164, previously identified as being required for the assembly of cytochrome *b*_6_*f* in *Arabidopsis* (Lennartz *et al*. 2001; Gabilly *et al*. 2010). *In vitro*, a recombinant form of CCS5 containing the WC*XX*C motif is redox-active and able to reduce a disulfide bond formed between the heme-linking cysteines of a soluble form of apocytochrome *f* (Lennartz *et al*. 2001; Gabilly *et al*. 2010). The cytochrome *c* assembly defect in the *ccs5*-null mutant can be chemically corrected by exogenously applied reducing agents, an observation demonstrating the physiological relevance of the disulfide reductase activity documented from the *in vitro* redox assays (Lennartz *et al*. 2001; Gabilly *et al*. 2010). Conversion of the disulfide-linked heme binding cysteines to thiols is also dependent upon CCDA whose activity is required to maintain the catalytic WC*XX*C motif in CCS5 in the reduced form (Page *et al*. 2004; Bushweller 2020). The proposed model is that the thiol-disulfide oxidoreductase CCDA and the thioredoxin-like protein CCS5/HCF164 define a trans-thylakoid pathway that sequentially transfers reducing power via a cascade of thiol-disulfide exchange from stroma to the oxidized C*XX*CH motif of apocytochrome *f* (Lennartz *et al*. 2001; Page *et al*. 2004; Motohashi and Hisabori 2006; Gabilly *et al*. 2010; Motohashi and Hisabori 2010; Gabilly *et al*. 2011). Maintenance of the C*XX*CH motif in the reduced form also relies on CCS4, a small protein predicted to be membrane-bound with a soluble domain facing the stroma (Gabilly *et al*. 2011). However, CCS4 contains no motif or residue suggestive of a redox biochemical activity, and orthologs can only be detected in a subset of green algae (Gabilly *et al*. 2011). The placement of CCS4 in the disulfide reduction pathway for plastid cytochrome *c* assembly was inferred from the finding that provision of reductants or the expression of an additional copy of *CCDA* suppresses the photosynthetic deficiency of the *ccs4-null* mutant (Gabilly *et al*. 2011). In this manuscript, we provide genetic and biochemical evidence for the involvement of CCS4 in two distinct pathways for reducing the disulfide-bonded heme-binding cysteines of apocytochrome *f* and discuss the possible functions of this assembly factor.

## MATERIALS AND METHODS

### Strains, media, and growth conditions

All algal strains used in this study are listed in Table S1 in supplemental material. Strains CC-124 and CC-4533 were used as wild-type; strains CC-4129, CC-5922 and ccs5-2 were used as *ccs5* mutants. The ccs5-2 strain is an insertional mutant (LMJRY0402.042908) from the CLiP library (Li *et al*. 2019) and carries the *APHVIII* cassette inserted in exon 3 of the *CCS5* gene (*Cre12*.*g493150*). The CC-4527 strain is the *ccs5* mutant (CC-4129) complemented with the wild type *CCS5* gene. This strain is referred to as ccs5(CCS5) and was described in Gabilly *et al*., 2010. Strains CC-4519, ccs4(pCB412) and CC-4525 were used as *ccs4* mutants. The ccs4ccs5(9) and ccs4ccs5(11) strains that displayed a tight photosynthetic deficiency had reverted at the end of the study. The suppressed strains were renamed SU9 and SU11 and deposited in the Chlamydomonas collection center as CC-5928 and CC-5929, respectively.

Wild-type and mutant *Chlamydomonas reinhardtii* strains were maintained on Tris-Acetate-Phosphate (TAP), with Hutner’s trace elements, 20 mM Tris-base and 17 mM acetic acid, or Tris-Acetate-Phosphate (TAP) supplemented with arginine (1.9 mM) (TARG) (Harris 1989). For some strains, TAP/TARG supplemented with 25 µg.ml^-1^ hygromycin B (TAP/TARG + Hyg) or 25 µg.ml^-1^ paromomycin (TAP/TARG + Pm) were used. All algal strains were grown on solid medium at 25°C in continuous light at 0.3 µmol/m^2^/s (Harris 1989). Solid medium contains 1.5% (w/v) agar (Select Agar_(tm)_, Invitrogen, 30391049).

Growth was tested on solid TAP/TARG, minimal medium, or minimal medium supplemented with 1.7 mM acetic acid under three different light conditions. For protein immunodetection, cells were grown in liquid TAP/TARG in an environmental incubator with continuous light at 0.3 µmol/m^2^/s and shaking at 180-190 rpm at 25°C.

Chemo-competent *Escherichia coli* DH5α strains were used as hosts for molecular cloning. The bacterial strains were grown at 37°C in Luria-Bertani (LB) broth and LB medium solidified with bacteriological agar (Silhavy *et al*. 1984). When antibiotic selection was required, 100 µg. ml^-1^ ampicillin was added.

### Genetic crosses

The experimental details for sexual crosses are described in former work (Subrahmanian *et al*. 2020a; Subrahmanian *et al*. 2020b). For molecular verification of the *ccs5* allele in the haploid progeny, the presence of the molecular tag (Phi “ϕ” flag) inserted in the *ARG7* intron used for insertional mutagenesis was detected via diagnostic PCR using the Phi-1 (5’-GTCAGATATGGACCTTGCT-3’) and Phi-2 (5’-CTTCTGCGTCATGGAAGCG-3’) primers (Gabilly *et al*. 2010). The presence of the wild-type *CCS4* allele, the *ccs4-1* mutation or suppressor mutations (*CCS4-2* and *CCS4-3*) was tested molecularly by diagnostic digestion of a PCR product. The alleles can be distinguished by virtue of a nucleotide polymorphism that abolishes the *Ban*II restriction site in the nonsense (*ccs4-1*) and missense mutations (*CCS4-2* and *CCS4-3*). The cell wall deficient strains CC-5925, CC-5922 and CC-5927 were generated by crossing the CC-4525 and CC-4517 strains and examining the progeny for the cell wall deficient trait, by resuspending cells in 0.1% Triton X-100. Untreated cells served as a control. Cells without a cell wall lyse in the presence of Triton X-100 (Hoffmann and Beck 2005). The *ccs4ccs5* mutants were generated by crossing the *ccs5* mutant (CC-4129) with the *ccs4* mutant (CC-4520) and selecting the progeny based on arginine prototrophy and paromomycin-resistance. For generating the vegetative diploids, the *ccs5::APHVIII* mutant from the CLiP collection was used. The CC-5923 and CC-5924 are haploid *ccs5::APHVIII arg7-8* progeny originating from the cross of the ccs5-2 strain by a wild type *arg7-8* strain (WT-PH2). WT-PH2 was obtained after four back-crosses of the CC-5611 (Subrahmanian *et al*. 2020a) by CC-4533 or CC-5155. The *ccs5*/*ccs5* diploids were constructed by crossing the CC-4129 strain by the CC-5924 strain and selection on TAP+Pm. The *ccs5* / *ccs5 CCS4-2* diploids were generated by crossing the CC-5931 strain by CC-4129 and selection on TAP+Pm. The CC-5931 strain is a haploid *ccs5::APHVIII CCS4-2 arg7-8* progeny issued from the cross of CC-5928 by CC-5923. The *ccs5*/*ccs5 CCS4-3* diploids were generated by crossing CC-5930 by CC-5924 and selection on TAP+Pm. The CC-5930 strain is a haploid *ccs5::ARG7*Φ *CCS4-3* progeny from the cross of CC-5929 by CC-5590. The wild-type diploids (+/+) were constructed by crossing CC-425 by CC-5590 and selection of the mitotic zygotes on TAP and already used in a former study (Barbieri *et al*. 2011).

### Growth assay and thiol-dependent rescue

Cells were grown on solid acetate-containing medium (TAP or TARG) for 5-7 days at 25°C under 0.3 µmol/m^2^/s illumination. Two loops of cells were resuspended in 500 µl water and optical density was measured spectrophotometrically at 750 nm and normalized to OD_750_ = 2. This normalized suspension was used as the starting material (1) to make five serial ten-fold dilutions (10^−1^, 10^−2^, 10^3^, 10^−4^, and 10^−5^). A volume of 5 µl from each dilution was plated on solid minimal medium (CO_2_) for photoautotrophic growth or TAP/TARG (acetate + CO_2_) for mixotrophic growth and incubated for 7 - 14 days under 30-50 µmol/m^2^/s illumination for growth assay. For thiol-dependent rescue, similar ten-fold serial dilutions were performed and equal volume from each dilution was plated on solid minimal medium (CO_2_) containing 1) either no MESNA (2-MercaptoEthane Sulfonate Na, Sigma, M1511-25G), or 2) 0.75 mM or 1.5 mM MESNA. Ten-fold serial dilutions were also plated on TAP/TARG (acetate + CO_2_) and incubated for 14 - 21 days under 30-50 µmol/m^2^/s for thiol-dependent rescue assay and under 0.3 µmol/m^2^/s with no exogenously added thiols as control.

### Fluorescence rise-and-decay kinetics

Fluorescence transients (also known as Kautsky effect) were measured using Handy FluorCam from Photon System Instruments that provides actinic illumination and saturating flash of light. Cells were grown mixotrophically on solid TAP/TARG under 0.3 µmol/m^2^/s illumination for 5-7 days at 25°C. To measure the fluorescence, cells are dark-adapted for 15 mins followed by a saturating flash of light. The fluorescence is recorded in arbitrary units (A.U.) over an illumination period of 5 seconds.

### Protein sample preparation

For heme staining and immunoblotting, two loops of cells grown on solid acetate-containing medium (TAP or TARG) for 5-7 days at 25°C under 0.3 µmol/m^2^/s illumination were collected and washed with 1-2 ml of 10 mM sodium phosphate buffer (NaH_2_PO_4_, pH 7.0), centrifuged at top speed and resuspended in 300 µl of the same buffer. Chlorophyll was extracted by mixing 10 µl of resuspended cells with 1 ml of acetone: methanol in a ratio of 80:20. The mixture was vortexed followed by centrifugation at 16,873 x *g* for 5 mins. Chlorophyll concentrations were then determined by measuring the optical density of the suspension at 652 nm. Concentrations for different strains were normalized to 1 mg/ml of chlorophyll, followed by cell lysis by two freeze-thaw cycles (samples were frozen −80°C for 1 hour and thawed in ice for ∼1 hour). Cells were then separated into membrane and soluble fractions by centrifugation for 5 mins at 16,873 x *g* (Howe and Merchant 1992). Membrane fractions corresponding to 10 µg of chlorophyll were separated electrophoretically by SDS-PAGE (12.5% acrylamide gel) for immunoblotting. For ECL-based heme staining, an amount of 10-20 µg of chlorophyll for each sample was used whereas for *in-gel* heme staining by TMBZ (3,3’,5,5’-tetramethylbenzidine, Sigma, 860336-1G), a quantity corresponding to 30-40 µg of chlorophyll per sample was loaded.

### Heme staining and immunoblotting

The presence of covalently bound heme in *c*-type cytochromes was detected by two methods, a) by enhanced chemiluminescence (ECL, Thermo Scientific, 34094) and b) *in-gel* TMBZ staining (3,3’,5,5’-Tetramethylbenzidine, Sigma, 860336-1G), both relying on the pseudo-peroxidase activity of heme (Wilks 2002). Detection of covalently bound heme in cytochrome *c* by ECL was done by separating proteins corresponding to 10-20 µg of chlorophyll on 12.5% acrylamide gel, followed by transfer to a PVDF membrane (Immobilon-P, IPVH00010) at 100 V (at 4°C) for 90 mins and treatment of the membrane with ECL reagent. *In-gel* heme staining was performed by soaking the gel in a solution of 6.3 mM TMBZ dissolved in methanol in the dark, mixed with 1 M sodium acetate for 1-1.5 hours in the dark with periodic shaking every 15-20 minutes. Bands revealing the heme peroxidase activity appeared immediately as a blue precipitate after addition of H_2_O_2_ (50% w/v) to a final concentration of 30 mM. For Immunoblotting, membrane fractions corresponding to 10 µg of chlorophyll were separated by SDS-PAGE and transferred as described above. Immunodetection was done by incubating the membrane with *Chlamydomonas* α-CF_1_ (1:10,000), α-PsbO (1:5000, Agrisera), α-cytochrome *f* (1:10,000), α-CCDA (1:1000), α-CCS5 (1:3,000). The CCDA and CCS5 antibodies were described in Motohashiand Hisabori, 2010 and Gabilly et al. 2010, respectively. The anti-cytochrome f antibody is produced against a recombinant form of an apocytochrome *f*-GST fusion. The α-rabbit (1:10,000) was used as secondary antibody.

### Construction of the suppressor strains

The *ccs4* strain expressing an additional copy of the *CCDA* gene was generated by introducing the construct pSL18-CCDA (Gabilly *et al*. 2011) in the CC-4525 strain via glass-bead method (Kindle 1990) and selecting the transformants on TARG+Pm plates. The suppressor strains were reconstructed by introducing constructs containing the *CCS4* gene from SU9 and SU11 into a Δ*ccs4ccs5* recipient strain. For comparison a *ccs5* strain was also reconstructed by introducing the wild-type *CCS4* gene in a Δ*ccs4ccs5* mutant. The *CCS4* gene (from ATG to stop including 165 bp of 3’ UTR) was cloned between the promoter of the beta-tubulin encoding gene (*TUB2*) and the *RBCS2* terminator in the pHyg3 plasmid (Berthold *et al*. 2002). The *CCS4* sequence from wild-type (CC-124), SU9 (CC-5928) and SU11 (CC-5929) was amplified with pHyg3-CCS4-F1 (5’-GTCACAACCCGCAAACGGGCCCATGTCGACCGGCATTGAG-3’) and pHyg3-CCS4-R1 (3’-ATTGTACTGAGAGTGCACCATATGCCAGCTACCTACAGTC-5’), which are *ApaI*- and *NdeI*-engineered primers using purified genomic DNA as a template. The digested PCR products were cloned at *Apa*I and *Nde*I sites in pHyg3 to yield the constructs pHyg3-CCS4(WT), pHyg3-CCS4(SU9) and pHyg3-CCS4(SU11), which were introduced into the CC-4518 strain by the glass bead transformation method (Kindle 1990). Transformants were selected on TAP+Hyg based on the Hyg resistance conferred by the iHyg3 cassette (containing the *APHVII* gene).

### Glass bead transformation

Recipient strains were cultured in liquid TAP or TARG and grown until exponential phase. At a cell density of 2-5 × 10^6^ cells/ml (measured by hemocytometer), cells were collected by centrifugation at 1,500 x *g* for 5 mins at 25°C, followed by resuspension and incubation in autolysin to a density of ∼1 × 10^8^ cells/ml for 45 – 90 mins (depending on the strain). The efficiency of enzymatic digestion of the cell wall was tested with 0.1% Triton X-100 as described above in the section detailing genetic crosses. Autolysin was removed by centrifugation of the treated cells at 1,500 x *g* for 5 mins at 25°C, and cells were resuspended in medium (TARG + 40 mM sucrose) to a final concentration of 1-2 × 10^8^ cells/ml. After autolysin treatment, 300 µl of cells, 100 µl of 20% PEG 8,000, 300 mg of glass beads and 1-2 µg of transforming DNA (linear or uncut) were mixed in a microfuge tube and vortexed for 30 – 45 secs at maximum speed. After glass beads settle, 300 – 350 µl of the mixture was transferred to 30 ml of TARG + 40 mM Sucrose, and incubated for 16-24 hrs at 25°C under 0.3 µmol/m^2^/s. After overnight recovery, cells are collected by centrifugation at 1,500 x *g* for 5 mins at 25°C, resuspended in 0.5 – 1 ml of TAP and plated on selective medium. Successful transformation yields colonies in 5-7 days.

### Bioinformatics

BLASTp, RPS-BLAST and PSI-BLAST searches (Altschul *et al*. 1997) against the non-redundant database at NCBI and BLASTp search against 1KP database (Matasci *et al*. 2014) were carried out with default parameters (1 February 2020). Protein domains were identified using Pfam (El-Gebali *et al*. 2019) and Conserved Domain (Marchler-Bauer *et al*. 2015) databases. Profile-profile searches for domain identification were carried out using CDvist (Adebali *et al*. 2015) and HHpred (SÖding *et al*. 2005) servers. Multiple sequence alignments were constructed using L-INS-I method in MAFFT program (Katoh *et al*. 2019). Sequence alignments were edited in Jalview (Waterhouse *et al*. 2009). Transmembrane regions and signal peptides prediction from amino acid sequences were carried out using Phobius (KÄll *et al*. 2004). Sequence logos were generated using Skylign (Wheeler *et al*. 2014).

## RESULTS

### CCS4 and CCS5 are functionally redundant

Previously, we showed that *CCS4* and *CCS5* are specifically required to maintain the heme-binding cysteines in the reduced form, a critical step for plastid cytochrome *c* assembly (Dreyfuss 1998; Xie *et al*. 1998; Gabilly *et al*. 2010; Gabilly *et al*. 2011). To define the contribution of each component in the assembly pathway, the *ccs4*-null (Δ*ccs4), ccs5*-null (Δ*ccs5)*, and *ccs4-*null *ccs5*-null double mutants (Δ*ccs4ccs5)* were first phenotypically assessed for photosynthetic growth, which requires a correctly assembled and functional cytochrome *f*, the membrane-bound plastid cytochrome *c* (Kuras *et al*. 1995; Zhou *et al*. 1996). To assess phototrophic growth, cells were grown under a light intensity of 30-50 µmol/m^2^/s, where they utilize atmospheric CO_2_ to perform photosynthesis. Compared to the Δ*ccs4* and Δ*ccs5* single mutants that show residual photosynthetic growth, the Δ*ccs4ccs5* double mutants are completely deficient for growth under phototrophic conditions (Figure 1A). Under low light (0.3 µmol/m^2^/s), all mutant strains can grow in the presence of acetate and atmospheric CO_2_ due to the dominant contribution of respiration *vs*. photosynthesis. Next, we performed chlorophyll fluorescence rise and decay kinetic, a non-invasive measurement of photosystem II (PSII) activity (Murchie and Lawson 2013). In wild type, the initial rise indicates electron movement through PSII, which is followed by a decay that indicates electron transfer through the cytochrome *b*_6_*f* (Figure 1B). When the energy absorbed by the chlorophyll in PSII cannot be utilized in the photosynthetic electron transport chain because of a complete block at cytochrome *b*_*6*_*f*, a saturating rise in fluorescence is observed with no decay phase thereafter. In Δ*ccs4ccs5*, the rise and plateau are a signature of a block at the level of cytochrome *b*_*6*_*f* complex due to the absence of holocytochrome *f* (Figure 1B). In Δ*ccs4* and Δ*ccs5*, the decay phase is less pronounced than in wild type, an indication that the strains retained cytochrome *b*_6_*f* functionality because some level of holocytochrome *f* is still assembled (Figure 1B), which agrees with the residual photosynthetic growth (Figure 1A). The level of holocytochrome *f* assembly in the different mutant strains was assessed biochemically via heme staining (Figure 1C). Holocytochrome *f* assembly is completely blocked in Δ*ccs4ccs5*, whereas the single null mutants accumulate partial levels of holocytochrome *f*, ∼10% in Δ*ccs5* and ∼>2% in Δ*ccs4*, respectively. Immunodetection of cytochrome *f* showed that the level of detected protein mirrors the level of heme revealed by heme staining (not shown). These results show that the synthetic photosynthetic growth defect of Δ*ccs4css5* is indeed due to a complete block in holocytochrome *f* assembly. The photosynthetic growth of the different mutant strains (Figure 1A) correlates with level of assembled holocytochrome *f* (Figure 1C). Loss of CCS4 function yields a more pronounced cytochrome *f* deficiency than loss of CCS5 function, whereas loss of both results in no cytochrome *f* assembly. Therefore, we concluded that CCS4 and CCS5 are functionally redundant for the provision of reductants for holocytochrome *f* assembly.

**Figure 1.**
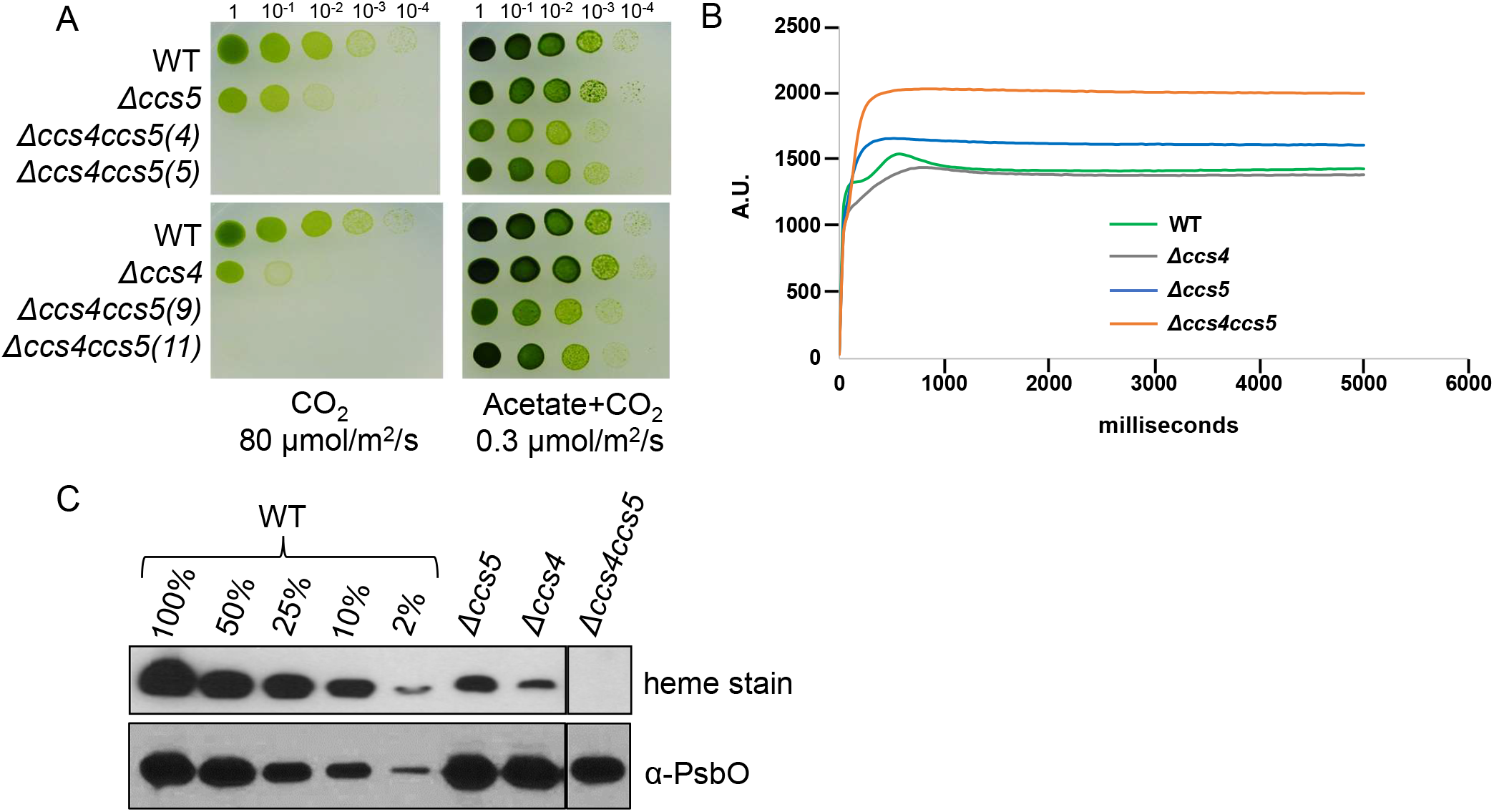
The Δ*ccs4ccs5* mutant exhibits a photosynthetic growth defect due to a complete loss of cytochrome *c* assembly. **(A)**. Ten-fold dilution series of wild type (WT, CC-124), *ccs5* (Δ*ccs5*, T78.15b-), *ccs4* (Δ*ccs4*, ccs4(pCB412)) and *ccs4ccs5* (Δ*ccs4ccs5*: ccs4 ccs5 (4), (5), (9), and (11)) strains were plated on acetate containing (right panel) and minimal (left panel) medium. Cells grown phototrophically (CO_2_) were incubated at 25°C for 7 days with 80 µmol/m^2^/s of light. Cells grown mixotrophically (acetate + CO_2_) were incubated at 25°C for 7 days with 0.3 µmol/m^2^/sec of light. **(B)** The fluorescence induction and decay kinetics observed in a dark-to-light transition of Δ*ccs4ccs5* is shown in comparison to that of Δ*ccs5*, Δ*ccs4*, and wild type. **(C)** Heme staining and α-PsbO immunodetection were performed on total protein extracts. Strains are as in 1A except for the Δ*ccs4* (*ccs4*.*1*) and Δ*ccs4ccs5* (ccs4ccs5(5)). Cells were grown mixotrophically on acetate with 0.3 µmol/m^2^/s of light at 25°C for 5-7 days. Samples correspond to 30 µg of chlorophyll. PsbO was immunodetected as loading control. The vertical line indicates cropping from the same gel.

### Loss of CCS4 and CCS5 can be chemically corrected by exogenous thiols

We previously evidenced that the partial deficiency in photosynthetic growth and holocytochrome *f* assembly in Δ*ccs4* and Δ*ccs5* mutants could be rescued by the application of exogenous thiols (Gabilly *et al*. 2010; Gabilly *et al*. 2011). Application of reducing agents such as MESNA (2-mercaptoethane sulfonate) or DTT (Dithiothreitol, not shown) can also partially rescue the Δ*ccs4ccs5* double mutant for phototrophic growth (Figure 2A). Using fluorescence rise-and-decay kinetics, we document that the chemical correction of the growth defect is due to a recovery of cytochrome *b*_6_*f* function, which indicates restoration of holocytochrome *f* assembly (Figure 2B). Note that a higher concentration of MESNA (4 mM *vs*.1.5 mM) was necessary to evidence a rescue of the Δ*ccs4ccs5* double mutant in the fluorescence transient experiments. We presumed that this difference is because cells are grown phototrophically under 30-50 µmol/m^2^/s for the growth assessment (Figure 2A) *vs*. mixotrophically under 0.3 µmol/m^2^/s for the measurement of fluorescence transient (Figure 2B). The Δ*petA* mutant, a null mutant in the plastid gene encoding apocytochrome *f* (Zhou *et al*. 1996), is completely blocked for photosynthetic growth and does not show any rescue with exogenous thiols, as expected.

**Figure 2.**
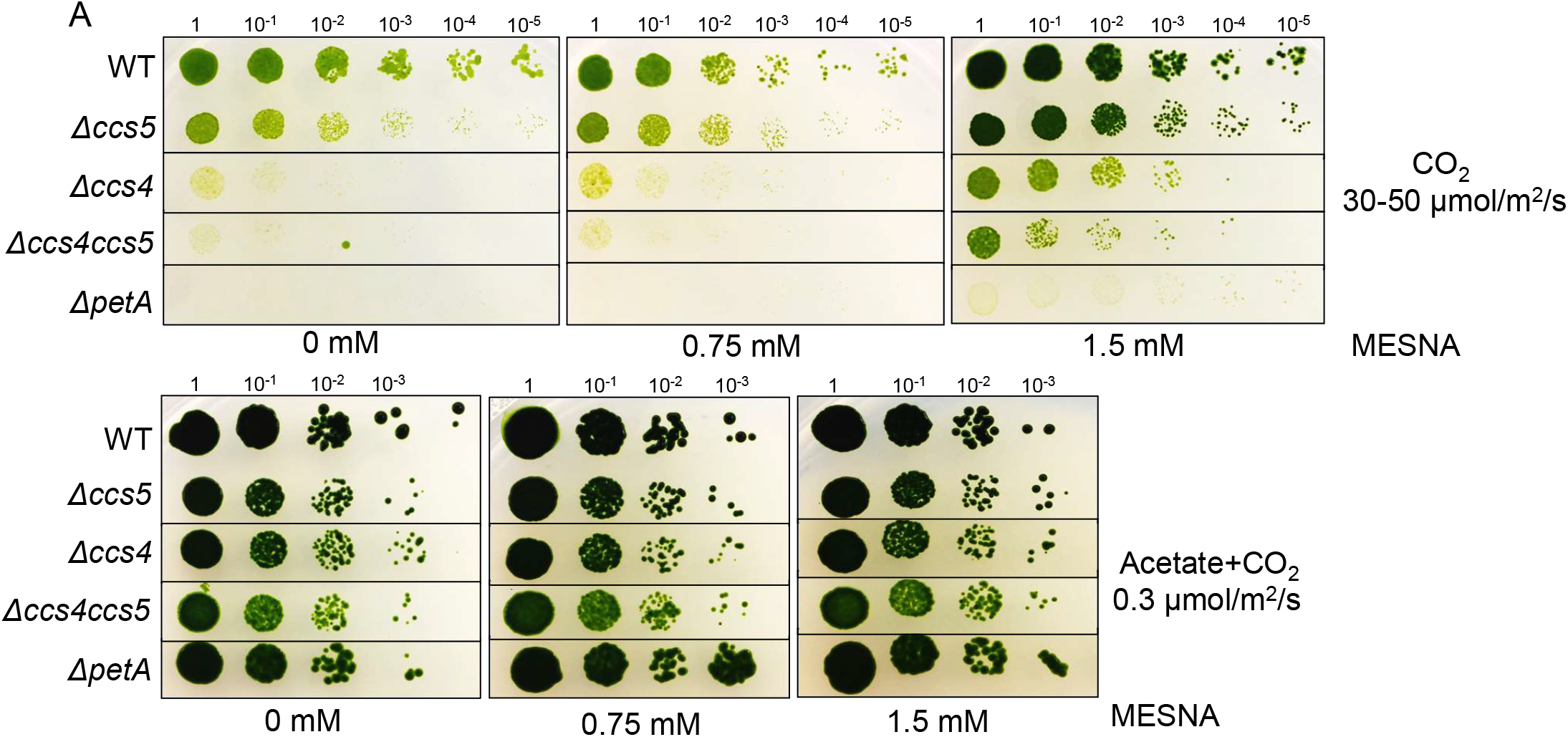
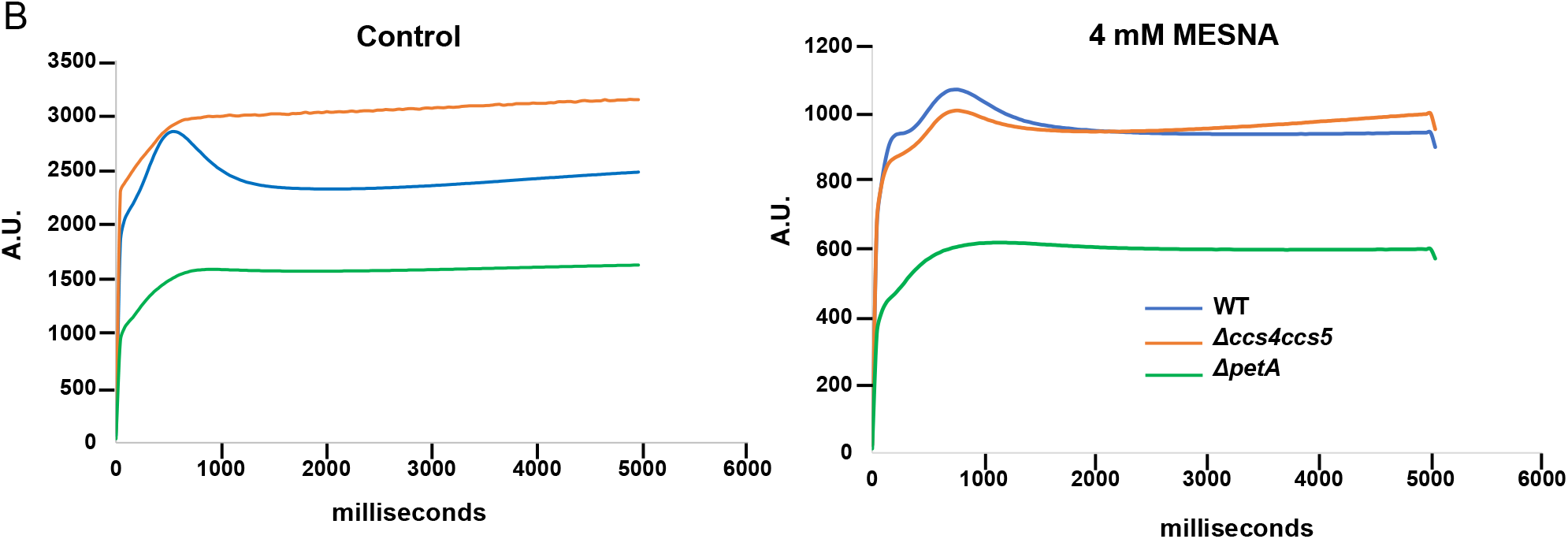
The photosynthetic defect in Δ*ccs4ccs5* is partially rescued by exogenous thiols. **(A)** Ten-fold dilution series of wild type (WT, CC-4533), *ccs5 (*Δ*ccs5*, ccs5(13)), *ccs4* (Δ*ccs4*, ccs4(28)), *ccs4ccs5* (Δ*ccs4ccs5*, ccs4 ccs5(35)) and Δ*petA* were plated on acetate and minimal medium with or without MESNA (2-mercaptoethane sulfonate sodium). Cells grown phototrophically (CO_2_) were incubated at 25°C for 20 days with 30-50 µmol/m^2^/s of light. Cells grown mixotrophically (acetate + CO_2_) were incubated at 25°C for 14 days with 0.3 µmol/m^2^/s of light. The horizontal lines indicate cropping from the same plate of serial dilution. **(B)** Fluorescence transients were measured on cells grown on solid acetate containing media for 5 days (with 0.3 µmol/m^2^/s of light) with or without MESNA..

### CCDA is down-accumulated in the *ccs4* mutant

Expression of an ectopic copy of *CCDA* encoding the trans-thylakoid thiol-disulfide oxidoreductase was shown to suppress Δ*ccs4* (Gabilly *et al*. 2011). This result led to the hypothesis that CCS4 might be involved in the stabilization of CCDA. To test this hypothesis, we monitored the abundance of CCDA by immunoblotting using an antibody against *Arabidopsis* CCDA but cross-reacting with the algal ortholog (Motohashi and Hisabori 2010). Because the original *CCDA* suppressed strain (Gabilly *et al*. 2011) lost the suppression over time, the strain was reconstructed. The suppression is weak but is best seen when the strain with an ectopic copy of *CCDA* is grown in mixotrophic conditions under 30-50 µmol/m^2^/s illumination (Figure 3A). The CCDA protein abundance decreased to ∼50% of the wild-type level in Δ*ccs4* but was restored to normal levels in the *CCS4* complemented strain and enhanced in the strain expressing an ectopic copy of the *CCDA* gene (Figure 3B). Conceivably, the decrease in CCDA abundance is not specific to the *ccs4* mutation and is a consequence of the loss of any CCS factor. That CCDA levels do not appear drastically changed in a *ccs5*-null mutant (compared to the corresponding complemented strain) indicates that the decrease in CCDA abundance is specific to loss of CCS4 function (Figure 3C). From this result, we concluded that CCDA down-accumulates when CCS4 function is compromised.

**Figure 3.**
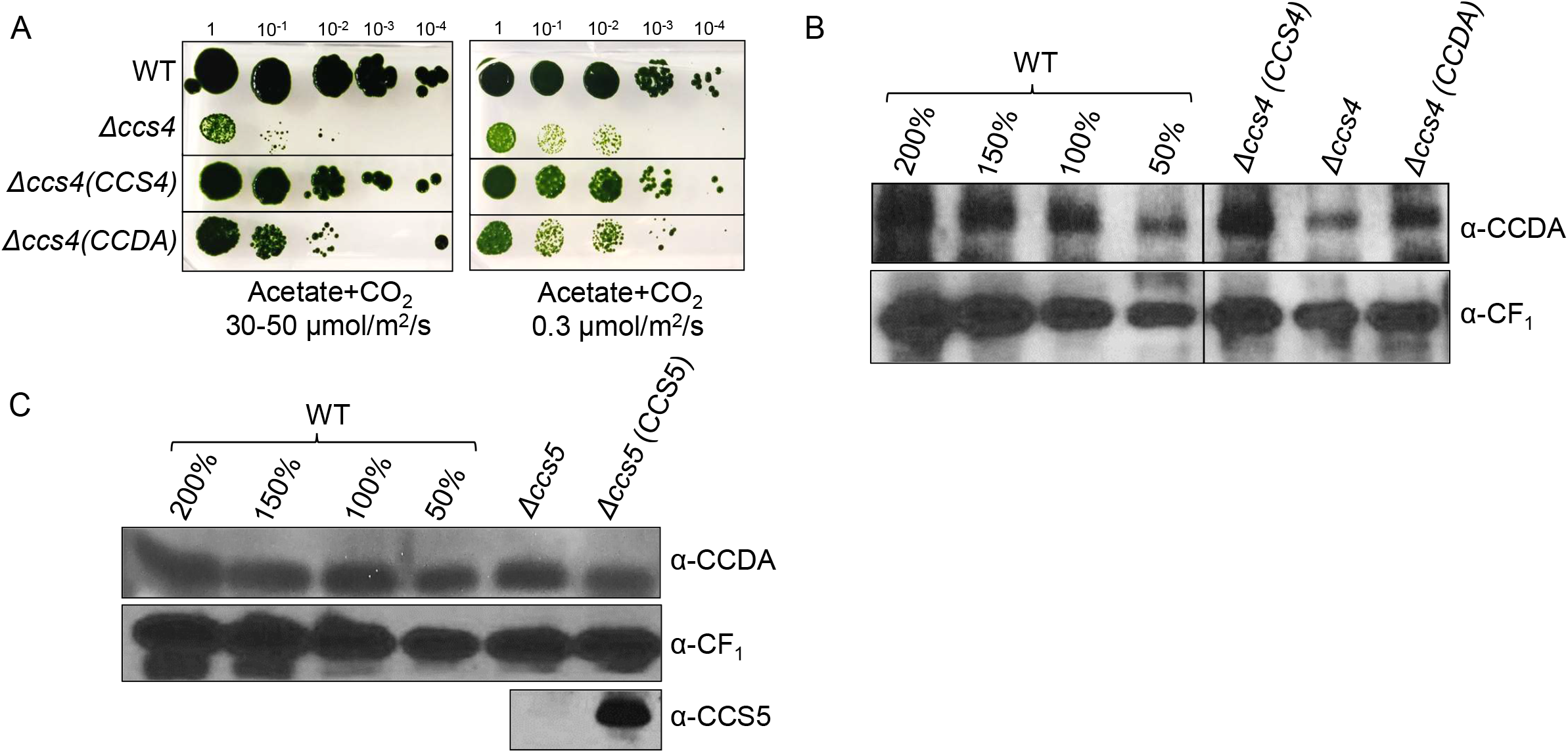
CCDA is downaccumulated in the *ccs4* mutant. **(A)** Suppression of the photosynthetic growth defect in the *ccs4* mutant by ectopic expression of *CCDA* was assessed by ten-fold dilution series. The wild type (WT, CC-124), *ccs4* (Δ*ccs4*, CC-4520), *CCS4* expressing (Δ*ccs4* (*CCS4*), CC-4522) and *CCDA* expressing (Δ*ccs4* (*CCDA*), CC-5926) *ccs4* strains were plated on acetate medium and grown mixotrophically (with 30-50 µmol/m^2^/s or 0.3 µmol/m^2^/s of light) at 25°C for 14 days. The horizontal lines indicate cropping from the same plate. **(B) and (C)** CCDA was immunodetected on total protein extracts of strains described in (a) and the *ccs5* (Δ*ccs5*, CC-4129) and corresponding complemented (Δ*ccs5*(*CCS5*), CC-4527) strains. Cells were grown mixotrophically (on acetate with 0.3 µmol/m^2^/s of light) at 25°C for 5-7 days. The vertical line indicates cropping from the same blot. Samples (100%) correspond to 10 µg of chlorophyll. Immunodetection with antisera raised against *Chlamydomonas* CF_1_ and CCS5 were used as control (Gabilly *et al*. 2010).

### Photosynthetic revertants of *ccs4ccs5* double mutant are restored for cytochrome *f* assembly

Over the course of this study, we noticed that two slow-growing Δ*ccs4ccs5* strains (#9 and #11, Figure 1A) had evolved to a faster growth phenotype. We suspected a possible reversion towards photosynthetic proficiency and named the strains SU9 and SU11 for suppressors of Δ*ccs4ccs5*#9 and #11, respectively. To test for the restoration of photosynthesis, we assessed the growth on acetate under high illumination. Under such conditions, cytochrome *b*_6_*f* deficient strains display significant photosensitivity (MalnoË *et al*. 2014). A Δ*ccs4ccs5* strain that had retained a tight photosynthetic deficiency due to loss of cytochrome *f* assembly displayed a photosensitive phenotype (Figure 4A). Interestingly, both SU9 and SU11 were resistant to high light treatment, suggesting the original Δ*ccs4ccs5*#9 and Δ*ccs4ccs5*#11 strains have recovered the *b*_6_*f* function (Figure 4A). Fluorescence transients confirmed restoration of the *b*_6_*f* function in SU9 and SU11 (Figure 4B). We evidenced that photosynthetic proficiency in SU9 and SU11 is due to recovery of holocytochrome *f* assembly, as shown by heme staining (Figure 4C). In SU9, cytochrome *f* assembly restoration is partial (∼25%) but in SU11, restoration appears to be wild type like (Figure 4C). The level of holocytochrome *f* assembly in SU9 and SU11 (Figure 4C) correlates with the recovery of photosynthetic proficiency assessed by growth in mixotrophic conditions and measurement of fluorescence transients (Figures 4AB). Immunodetection of cytochrome *f* showed that the level of detected protein mirrors the level of heme revealed by heme staining (Fig. S1).

**Figure 4.**
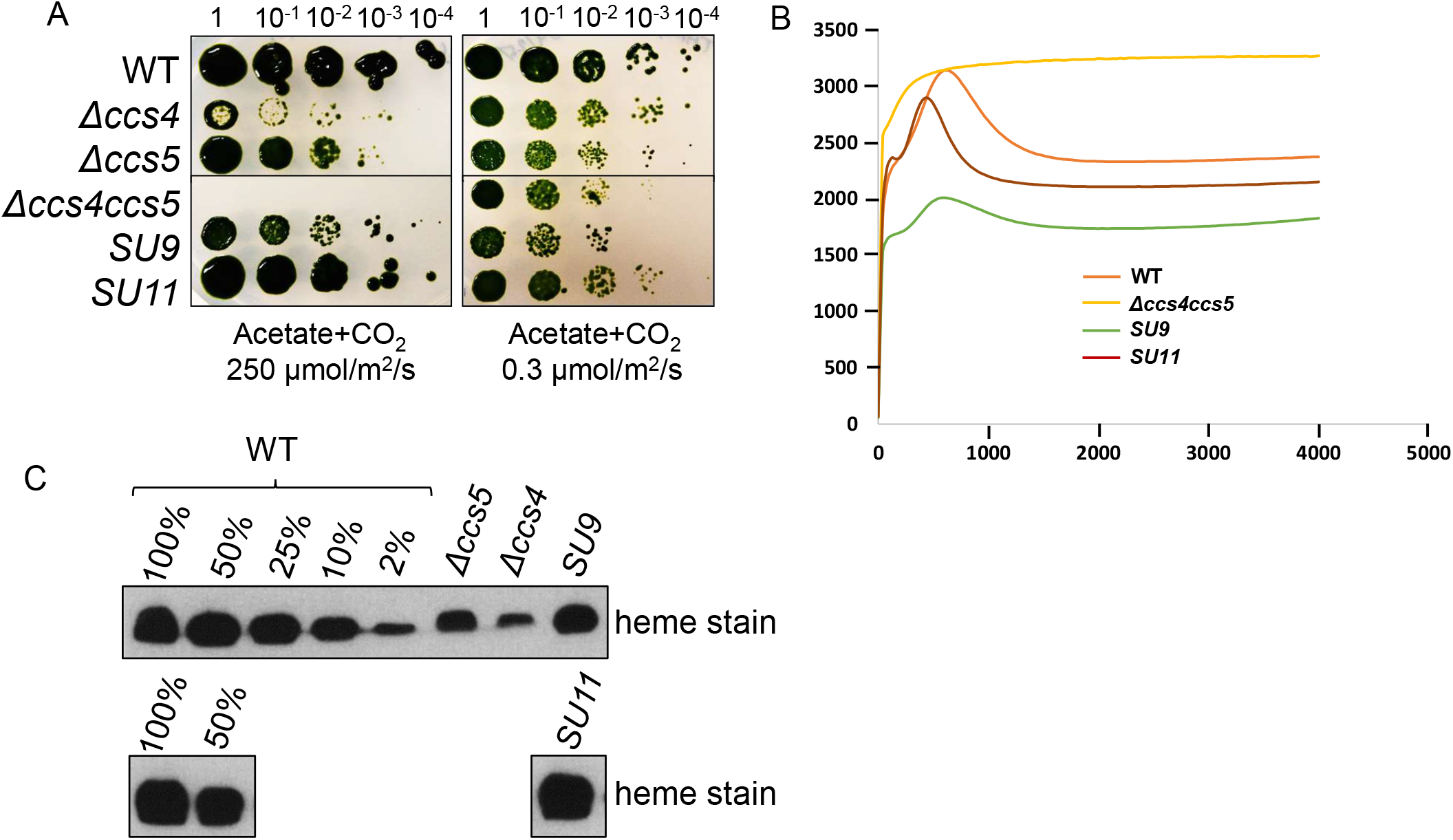
Suppressors of Δ*ccs4ccs5* are restored for phototrophic growth and cytochrome *f* assembly. **(A)** Ten-fold dilution series of wild type (WT, CC-124), *ccs5* (Δ*ccs5*, T78.15b-), *ccs4* (Δ*ccs4*, ccs4-1) and *ccs4ccs5* (Δ*ccs4ccs5*: ccs4 ccs5 (5)), SU9 and SU11 were plated on acetate medium and grown mixotrophically (with 250 µmol/m^2^/s or 0.3 µmol/m^2^/s of light) at 25°C for 14-18 days. The horizontal line indicates cropping from the same plate of serial dilutions. **(B)** Heme staining and α-PsbO immunoblotting were performed on total protein extracts prepared from cells (same strains as in A) grown mixotrophically (on acetate with 0.3 µmol/m^2^/s of light) at 25°C for 5-7 days. Samples (100%) correspond to 30 µg of chlorophyll. Immunodetection with antisera against PsbO was used as loading control. **(C)** The fluorescence induction and decay kinetics observed in a dark-to-light transition of the SU9 and SU11 suppressors are shown in comparison to Δ*ccs4ccs5* and wild type. Strains are the same as in A.

### Reversion of the *ccs4ccs5* is due to spontaneous dominant gain-of-function mutations in *CCS4*

Next we sought to determine the molecular basis of the reversion in SU9 and SU11. We ruled out the possibility that a suppressor mutation maps to the *CCS5* gene, considering a 3 kb region containing the entire coding sequence of *CCS5* is missing in Δ*ccs5* (Gabilly *et al*. 2010). Instead, we suspected that the point mutation (C to T) at a codon (**C**AG) encoding a glutamine (Q_50_) in the *CCS4* gene might have reverted in SU9 and SU11 (Gabilly *et al*. 2011). Sequencing of the *CCS4* gene in SU9 revealed the nonsense mutation in the original *ccs4* mutant (TAG) was changed to a missense mutation (T**G**G) resulting in a tryptophan (W) residue (Figure 5A). Surprisingly, in SU11, the *CCS4* gene carried an in-frame deletion of 3 codons removing the stop codon (GCT**TAG**ATG), resulting in a CCS4 protein shorter by 3 residues (Figure 5A). The intragenic mutations in the *CCS4* gene alter residues in the soluble domain of CCS4, which is predicted to face the stromal side of the thylakoid membrane (Gabilly *et al*. 2011).

**Figure 5.**
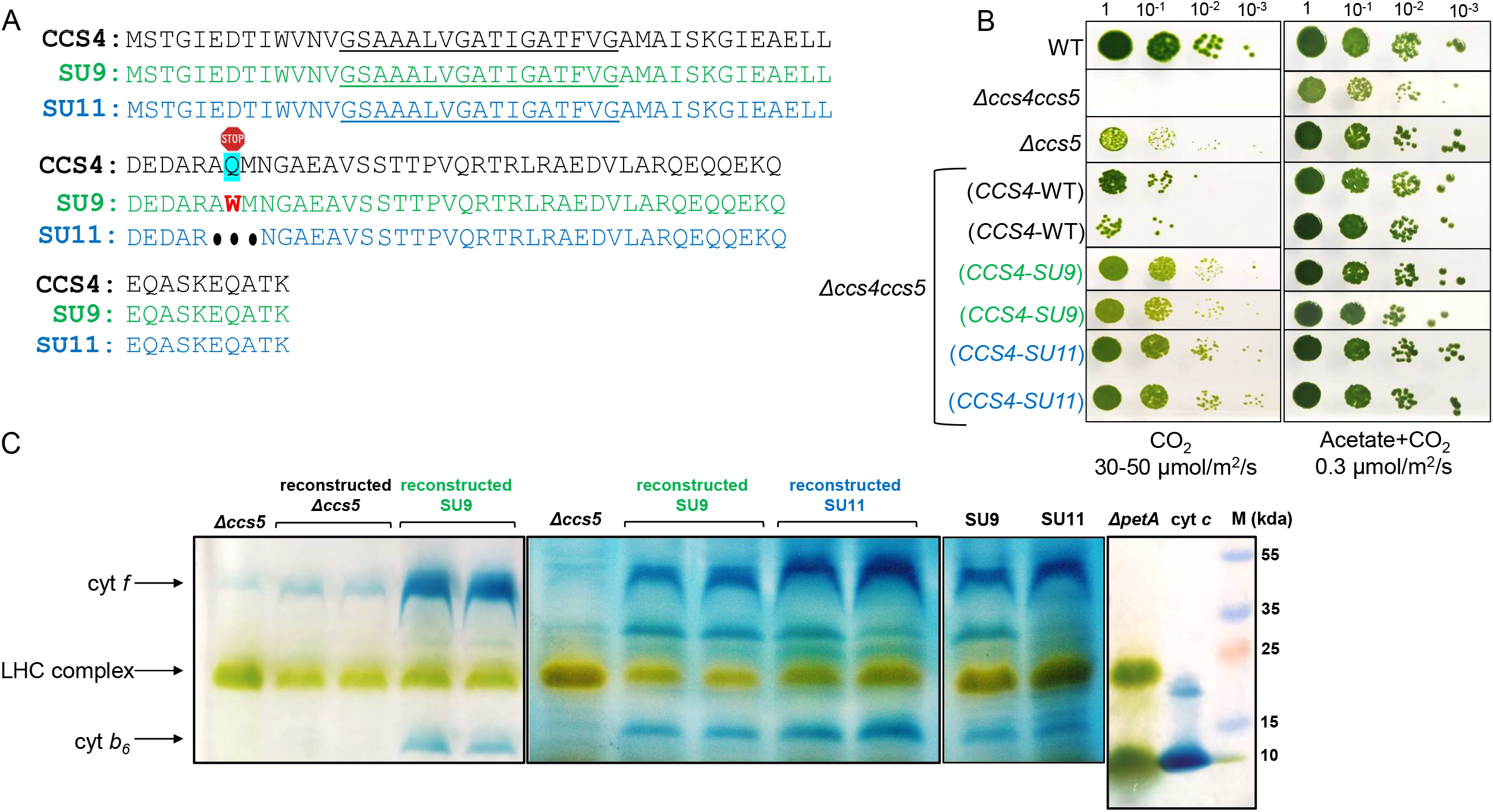
Mutations in *CCS4* suppress the cytochrome *f* assembly defect in the *ccs5* mutant. **(A)** Amino acid sequence of CCS4 deduced from the *CCS4* gene in the original *ccs4-1* mutant, SU9 and SU11 are shown in black, green, and blue, respectively. The predicted transmembrane domain is underlined. The putative transmembrane domain was predicted using the TMHMM-2.0 prediction tool (Krogh *et al*. 2001). Sequence analysis of the *CCS4* gene shows that the original nonsense mutation (resulting in stop at codon 50 encoding for glutamine) is now a missense mutation (resulting in the codon 50 encoding tryptophan) in SU9 and an in-frame deletion mutation (resulting in a deletion of the three residues A_49_Q_50_M_51_) in SU11. **(B)** Wild-type (WT, CC-124), *ccs5* (Δ*ccs5*, T78.15b-), *ccs4ccs5* (Δ*ccs4ccs5*, ccs4 ccs5 (5)) and the reconstructed *ccs5* (CCS4-WT), SU9 (CCS4-SU9), and SU11 (CCS4-SU11) strains were used for ten-fold dilution series. The Δ*ccs5*, SU9 and SU11 were reconstructed by introducing the constructs pHyg3-CCS4(WT), pHyg3-CCS4(SU9) and pHyg3-CCS4(SU11) into a Δ*ccs4ccs5* recipient strain. Two independent transformants for each construct are shown. Cells were plated on minimal medium and grown phototrophically (with 30-50 µmol/m^2^/s of light) or mixotrophically (with 0.3 µmol/m^2^/s of light) at 25°C for 14-18 days. The horizontal lines indicate cropping from the same plate of serial dilutions. **(C)** *In-gel* heme staining was performed on total protein extracts corresponding to 30 µg of chlorophyll. Samples from the Δ*ccs5* and original SU9 and SU11 were loaded on the gel for comparison to the reconstructed strains. The strains are *ccs5* (Δ*ccs5*, T78.15b-), the reconstructed *ccs5* (CCS4-WT), SU9 (CCS4-SU9), SU11 (CCS4-SU11) and Δ*petA*. The level of LHC complex is used as a loading control. One µg equine heart cytochrome *c* (∼10 kDa; Sigma) is used as a control for the peroxidase activity.

To ascertain that the molecular changes identified in *CCS4* were responsible for the suppression, we performed genetic analysis by crossing SU9 and SU11 to the original Δ*ccs4ccs5*, or Δ*ccs4* or Δ*ccs5* mutants. However, despite several attempts, no complete tetrads and very few viable haploid progenies were recovered from our genetic crosses. We opted to reconstruct the suppressed strains in the background of the Δ*ccs4ccs5* strain. The *CCS4* gene from wild-type, SU9 and SU11 strains was cloned in a plasmid and the resulting constructs were introduced into a Δ*ccs4ccs5* mutant followed by analysis of photosynthetic growth and holocytochrome *f* assembly from representative transformants. In comparison to the recipient Δ*ccs4ccs5*, (that is completely blocked for photosynthetic growth), transformants expressing the wild-type *CCS4* gene showed the phenotype of the Δ*ccs5* strain, whereas transformants expressing *CCS4* with the mutations exhibit enhanced photosynthetic growth (Figure 5B). The phenotypes of the reconstructed suppressed strains mirrored that of the original SU9 and SU11, with the reconstructed SU11 displaying a greater suppression of the photosynthetic defect than reconstructed SU9. We further demonstrated that the restoration of photosynthetic growth in the reconstructed suppressed strains is due to restoration of cytochrome *f* assembly, as shown by heme staining (Figure 5C). We also noted the restoration of holocytochrome *b*_6_ accumulation, an indication that cytochrome *b*_6_*f* complex assembly is also restored (Figure 5C). In both the original SU9 and SU11 and the reconstructed suppressed strains, the mutations in *CCS4* restore the level of holocytochrome *f* assembly above that of the *ccs5*-null mutant, an indication that these mutations confer a gain-of-function. The photosynthetic growth of *ccs5* x SU9 and *ccs5* x SU11 diploids is enhanced compared to that of *ccs5* x *ccs5* diploids (Figure S2). This indicates that the *CCS4* mutations in SU9 and SU11 are dominant, as expected for gain-of-function mutations. We concluded that dominant suppressor mutations in *CCS4* are the molecular basis for the restoration of cytochrome *f* assembly in the SU9 and SU11 strains. We named the suppressor mutations in SU9 and SU11, *CCS4-2* and *CCS4-3*, respectively.

### CCS4 – like proteins occur in the green lineage

With the exception of orthologs detected in close relatives of *Chlamydomonas*, we previously reported that CCS4 does not appear to be evolutionarily conserved (Gabilly *et al*. 2011). CCS4-like proteins in algal species related to *Chlamydomonas* are small proteins (<100 residues) with a putative transmembrane domain suggesting anchoring to the membrane (Figure S3). Remarkably, the C-terminal domain of algal CCS4 orthologs is predicted to face the stroma and is characterized by an abundance of charged residues (Figure S3). To identify CCS4-like proteins that might have considerably diverged, we performed an exhaustive search using PSI-BLAST (Data S1). CCS4-like proteins were identified in several species of green algae and land plants, which constitute the *viridiplantae* or entire green lineage (Data S1, Dataset 1A, Figure 6A). Interestingly, our analysis also revealed that CCS4 orthologs are related to COX16, a protein involved in the assembly of cytochrome *c* oxidase, a respiratory enzyme required for energy-transduction in mitochondria (Carlson *et al*. 2003) (Data S1, Dataset 1B and Figure 6B). First identified in yeast, COX16 is a mitochondrial inner membrane bound protein with a C-terminal domain facing the intermembrane space (Carlson *et al*. 2003). The biochemical activity of COX16 still remains enigmatic but its placement in a disulfide reducing pathway required for copper incorporation into the Cox2 subunit of cytochrome *c* oxidase suggests COX16 and CCS4 might share some commonalities in their biochemical activity (Su and Tzagoloff 2017; Aich *et al*. 2018; Cerqua *et al*. 2018).

**Figure 6.**
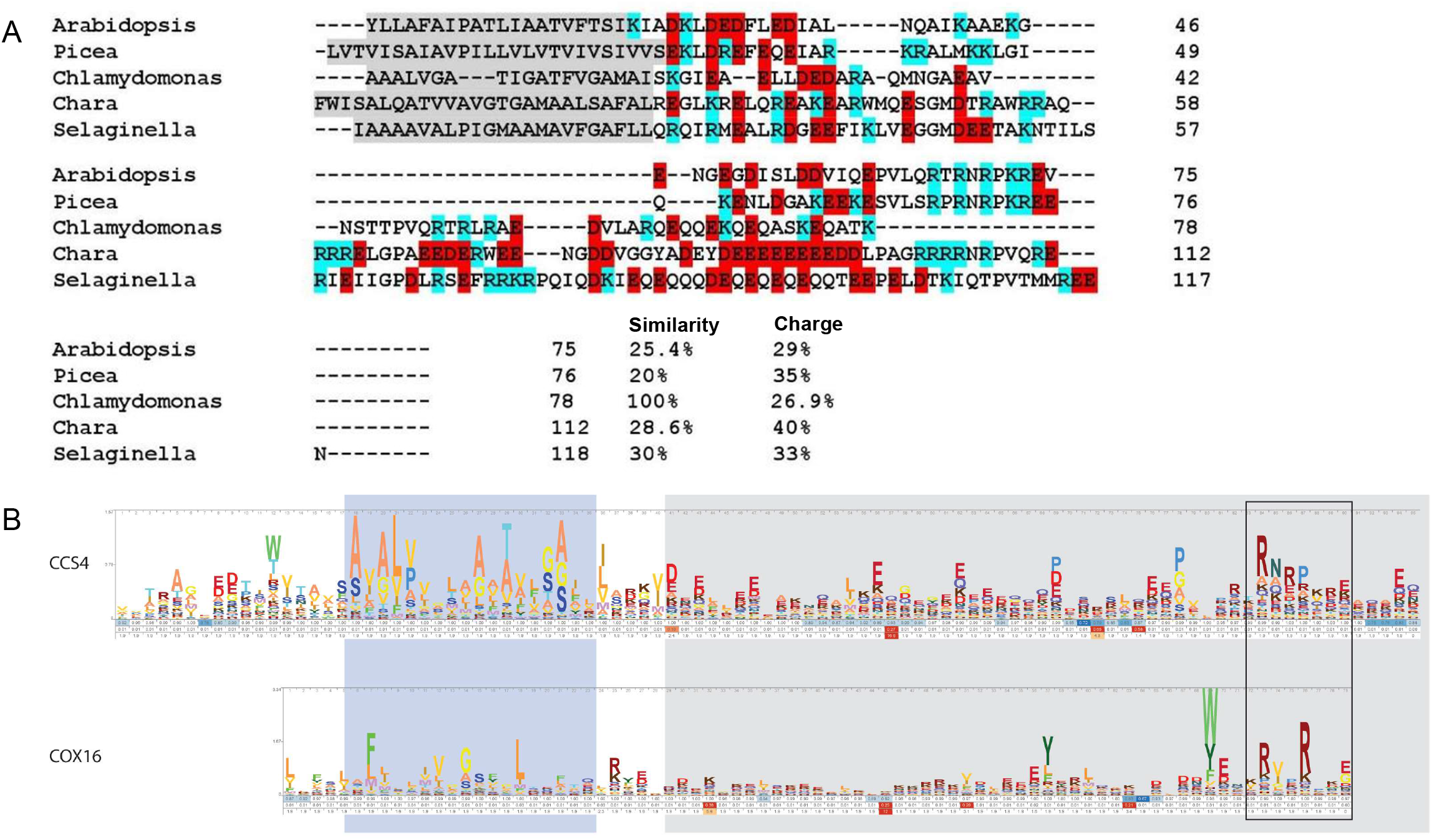
CCS4-like proteins are present throughout the green lineage. **(A)** Sequences of *Arabidopsis thaliana* (accession no. NP_194884.1), *Picea sitchensis* (accession no. ABK22505.1), *Chlamydomonas reinhardtii* (accession no. ADL27744.1), *Chara braunii* (accession no. GBG73810.1), *Selaginella moellendorffii* (accession no. XP_024543966.1) were aligned using Clustal omega (Sievers and Higgins 2014) and manually edited in Word. The transmembrane domains are highlighted in grey, residues with positive and negative charge are highlighted in blue and red, respectively. **(B)** Sequence logo comparison of CCS4 and COX16. The blue background indicates transmembrane domains, the grey background indicates charged amino acid rich regions and the rectangle highlights a similar C-terminal motif.

## DISCUSSION

### CCS4 and CCS5 are functionally redundant in the delivery of reductants

In all bacteria and plastids, cytochrome *c* maturation relies on the transfer of reducing power from the *n*-side to the *p*-side via sequential thiol-disulfide exchanges involving a membrane thiol-disulfide oxidoreductase of the DsbD family and a thioredoxin-like protein (Gabilly and Hamel 2017; Bushweller 2020; Das and Hamel 2021). Operation of this transmembrane pathway is critical to counter the oxidation of the heme-linking cysteines in the C*XX*CH motif of apocytochrome *c* into a disulfide (Gabilly and Hamel 2017; Das and Hamel 2021). In *Arabidopsis*, loss of CCDA, the thylakoid membrane thiol-disulfide oxidoreductase of the DsbD family or the thioredoxin-like protein CCS5/HCF164 results in a partial cytochrome *f* deficiency (Lennartz et al. 2001; Page et al. 2004). Because there is no other CCDA and CCS5/HCF164 encoding genes in *Arabidopsis*, this result was taken as an indication that a CCDA-independent route for the provision of reductants must exist (Lennartz et al. 2001; Page et al. 2004). In bacteria, loss of the membrane thiol-disulfide oxidoreductase or the thioredoxin-like protein results in a complete block in cytochrome *c* maturation, suggesting that redundancy in the provision of reductants is specific to the assembly of *c*-type cytochromes in the plastid (Beckman and Kranz 1993; SchiÖtt *et al*. 1997; Fabianek *et al*. 1998; Beckett *et al*. 2000; Deshmukh *et al*. 2000). In *Chlamydomonas*, the *ccs4*-null and *ccs5*-null mutants are still able to grow photosynthetically because they are partially deficient in plastid holocytochrome *f* (Gabilly et al. 2010; Gabilly et al. 2011). However, the *ccs4*-null *ccs5*-null double mutant displays a synthetic photosynthesis-minus phenotype characterized by a complete loss of holocytochrome *f* accumulation, an indication that CCS4 and CCS5 are redundant (Figure 1). Evidence that Δ*ccs4ccs5* can be rescued by application of exogenous thiols similarly to each single Δ*ccs4* and Δ*ccs5* mutants (Gabilly *et al*. 2010; Gabilly *et al*. 2011 and Figure 2) solidifies the view that the complete block in plastid cytochrome *c* maturation is attributed to a defect in the provision of reductants.

### CCS4 is required to stabilize CCDA

The finding that ectopic expression of *CCDA* partially suppresses the photosynthetic deficiency of the *ccs4* mutant led to the proposal that CCS4 might function in the reducing pathway by stabilizing CCDA (Gabilly et al. 2011). CCDA levels are significantly decreased due to loss of CCS4 function and restored upon complementation with the *CCS4* gene or suppression by *CCDA* (Figure 3). Because the *CCDA* gene is not downregulated in the *ccs4* mutant (Gabilly *et al*. 2011), a plausible scenario is that CCDA is still synthesized but destabilized in the absence of CCS4 and presumably partially turned over (Figure 3). One possibility is that the transmembrane domain of CCS4 is the structural element required for stabilizing the CCDA protein. However, a singular function of CCS4 as a stabilizer of CCDA is probably an oversimplification. The fact that loss of CCS4 yields a more drastic phenotype than loss of CCS5 speaks to an additional activity for CCS4 beyond its function of stabilizing CCDA. Moreover, the synthetic phenotype exhibited by Δ*ccs4ccs5* suggests that CCS4 and CCS5 are redundant and operate in different reducing pathways (Figures 1 and 4).

### CCS4 is a redox component of CCS5-independent pathway

Evidence for the involvement of CCS4 in a CCS5-independent reducing pathway was further substantiated with the isolation of dominant *CCS4-2* and *CCS4-3* mutations in the photosynthetic revertants of the Δ*ccs4ccs5* mutants SU9 and SU11, respectively (Figures 4 and 5). In the *CCS4-2* mutation, the original stop codon was changed to a codon encoding a tryptophan (instead of a glutamine in the wild-type *CCS4*) while in the *CCS4-3* mutation, an in-frame deletion produces a CCS4 protein which is shorter by three amino-acids (Figure 5A). Suppressor mutations in the *CCS4* gene restore cytochrome *f* accumulation above the residual levels in the *ccs5* mutant, suggesting that these are gain-of-function mutations (Figures 4 and 5). The mechanism of suppression is unclear but one possible interpretation is that alteration of the stroma-facing C-terminal domain of CCS4, where the changes occur, enhance the delivery of reductants to the lumen when the CCDA/CCS5 pathway no longer operates. Indeed, the *CCS5* gene is deleted in SU9 and SU11 and hence, the CCDA/CCS5 pathway is presumably no longer functional. We can imagine that in SU9 and SU11, delivery of reducing power to the lumen is “boosted” due to the gain-of-function mutations in *CCS4*, uncovering the role of CCS4 in a CCDA/CCS5 independent pathway Whether CCDA is also required for the CCS5-independent pathway remains an open question as there is currently no available *ccdA* mutant in *Chlamydomonas* to test this hypothesis.

### Operation of two trans-thylakoid disulfide reducing pathways

We propose that maintenance of the heme-binding cysteines in the reduced state in the lumen depends on two trans-thylakoid disulfide reducing pathways (see graphical abstract). One pathway, conserved in bacteria and referred to as the CCS5-dependent pathway (pathway 1), relies on the activity of CCDA that provides reductants to CCS5/HCF164, and another pathway is CCS5-independent (pathway 2). In pathway 1, CCS4 stabilizes CCDA and is also required for the CCDA-dependent transfer of reducing power, possibly by recruiting reductant to CCDA. CCS4 is also a component of the second pathway that controls the provision of reductant for the heme attachment reaction, named pathway 2 (Graphical abstract). Based on our understanding of the process, the pathway 2 must operate with a) a reductant originating from the stroma and transferred across the thylakoid membrane to the lumen. If CCDA is required for pathway 2, the source of reducing power must be transduced via thiol-disulfide exchanges from stroma to lumen. If CCDA is not required for this pathway, a distinct mechanism for transferring reducing power across must exist as there is no other CCDA-like protein in photosynthetic eukaryotes. Regardless of the involvement of CCDA in the CCS5—independent pathway, one or several lumen resident enzyme(s) must utilize a form of reducing power to maintain the apocytochrome *f* heme binding cysteines under the reduced form. The identity of such disulfide reductase enzymes remains unknown. If the operation of the transmembrane pathways is critical to counter the oxidation of the heme-linking cysteines in the CXXCH motif into a disulfide, the nature of the oxidant is unknown (Gabilly and Hamel 2017; Das and Hamel 2021). In bacteria, there is genetic evidence that the heme-linking cysteines are first targets of the disulfide bond forming machinery and subsequently reduced via the transmembrane disulfide reducing pathway prior to the heme ligation reaction (Erlendsson and Hederstedt 2002; Erlendsson et al. 2003; Turkarslan et al. 2008; Small et al. 2013). We envision that a similar mechanism exists in the thylakoid lumen. In *Arabidopsis*, a disulfide bond forming enzyme named LTO1 (Lumen Thiol Oxidoreductase1) has been described to introduce a disulfide bond in PsbO, a PSII subunit (Feng *et al*. 2011; Karamoko *et al*. 2011). That the CXXCH of apocytochrome *f* is also a relevant target of action of LTO1 still awaits experimental testing.

### Conservation in the green lineage and evolutionary relationship to COX16

In this study, we also provide evidence for the occurrence of highly divergent CCS4-like proteins in *viridiplantae*, underscoring the conservation of CCS4 function in plastid cytochrome *c* maturation. Interestingly, HCF153, the CCS4-like protein in *Arabidopsis* (Figure 6) localizes to the plastid and is tightly bound to the thylakoid membrane (Lennartz *et al*. 2006). A possible function in holocytochrome *f* maturation is supported by the fact that loss of HCF153 elicits a cytochrome *b*_6_*f* assembly defect (Lennartz *et al*. 2006). That HCF153 is the functional equivalent to CCS4 is very likely but remains to be demonstrated. CCS4-like proteins in the plastid are related to COX16-like proteins, which exhibit a transmembrane domain and a charged C-terminal moiety (Figure 6B). Considering there are no residues or motifs speaking to a reducing activity, CCS4 and COX16 might function in a disulfide reducing pathway to recruit the source of reducing power or create a microenvironment for efficient use of the reductant. In plastids, stromal thioredoxin-*m* is the proposed source of electrons to CCDA based on *in organello* experiments (Motohashi and Hisabori 2010). A similar function can be envisioned for COX16 but the source of electrons in the disulfide reducing pathway for metalation of Cox2 is currently unknown (Swaminathan and Gohil 2022).

## DATA AVAILABILITY

Strains and plasmids used in this study can be requested at the *Chlamydomonas* collection center. The authors affirm that all data necessary for confirming the conclusions of the article are present within the article, figures, and tables.

## ACKNOWLEDGMENTS

This work was supported by a U.S. Department of Energy (DOE), Office of Science, Basic Energy Sciences (BES) grant (DE-SC0014562) to P.H. and grant R35GM131760 from National Institutes of Health (to I.B.Z.)

---

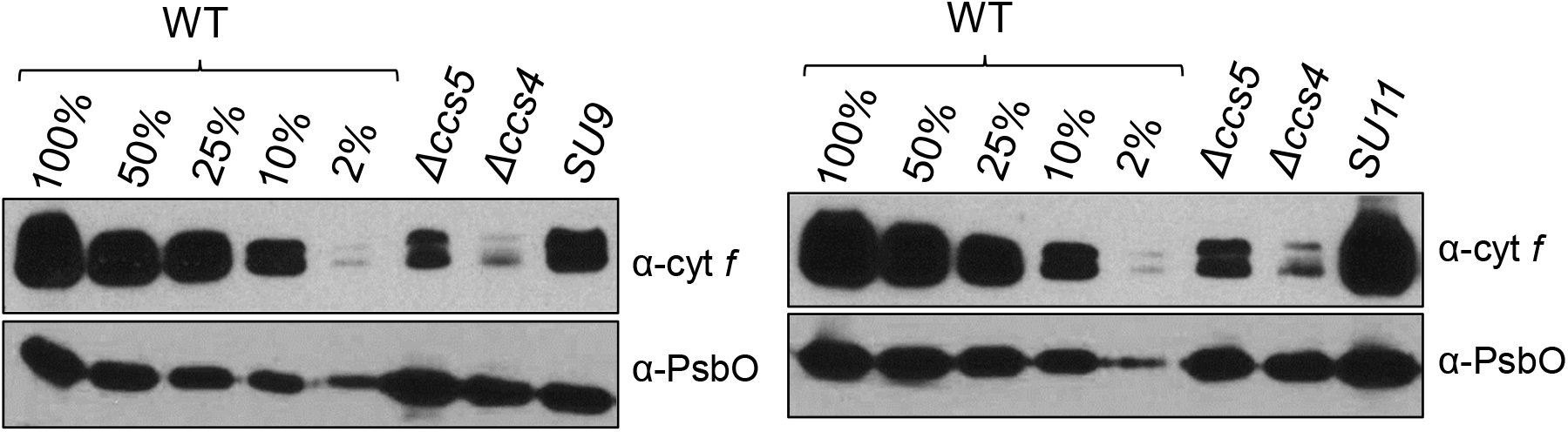

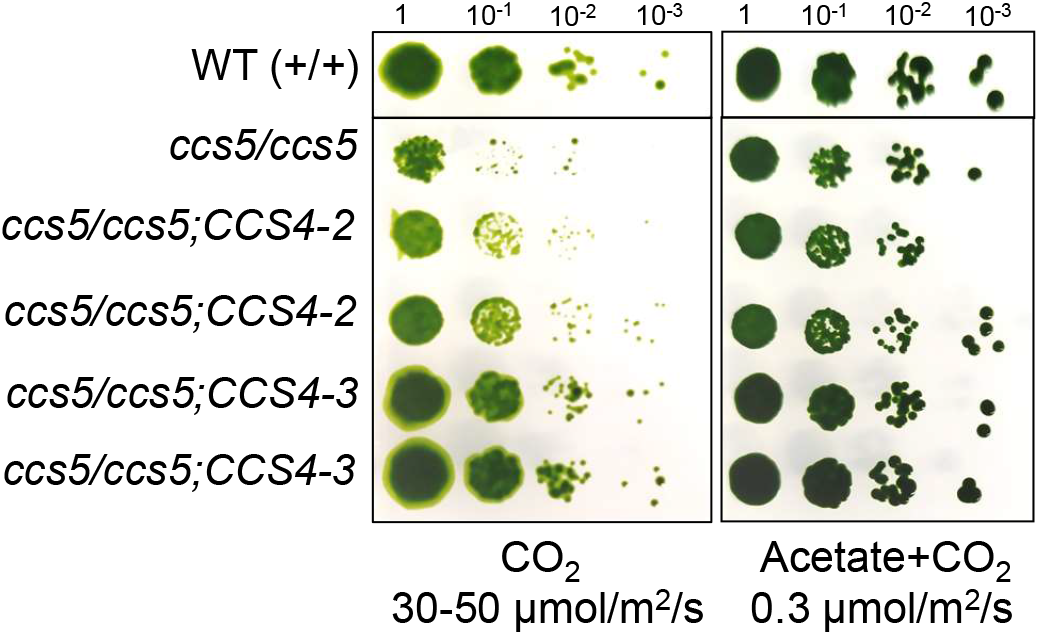

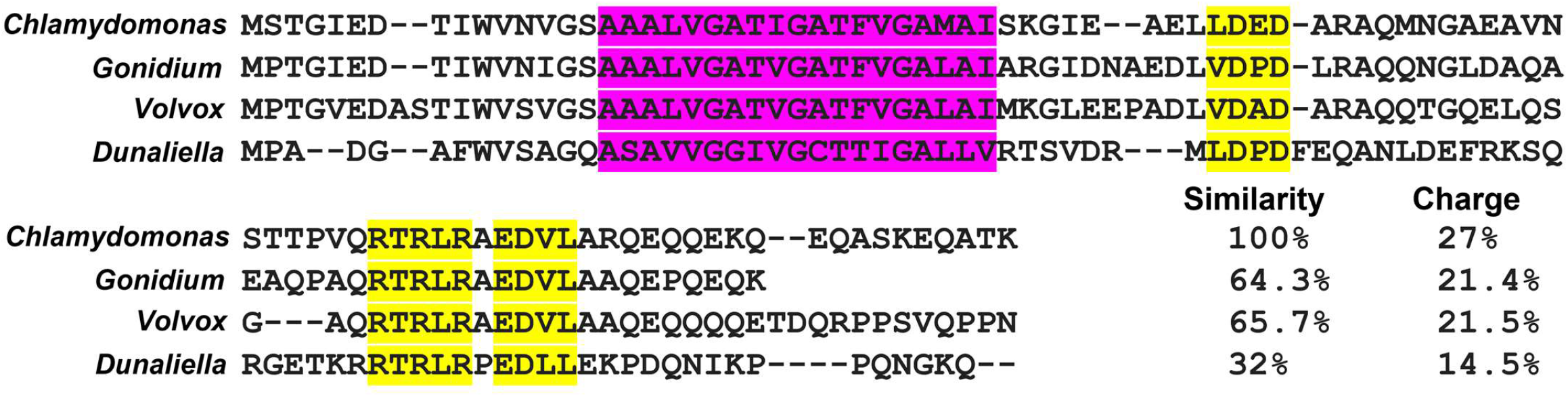

