## Supplementary material for "TWO DISULFIDE-REDUCING PATHWAYS ARE REQUIRED FOR THE MATURATION OF PLASTID *C*-TYPE CYTOCHROMES IN *CHLAMYDOMONAS REINHARDTII*": Dataset S1

Dataset S1. Multiple sequence alignments.

Dataset S1A. CCS4 homologs from a representative set of the *Viridiplantae* clade. Transmembrane regions are highlighted in grey; similar charged amino acids motifs are colored in red.

```
KXZ48069.1_hypothetical_protein_GPECTOR_30g164_[Gonium_pectorale]                      MP----------------------------------------------------------
ADL27744.1_cytochrome_c_synthesis_4_protein_[Chlamydomonas_reinhardtii]                MS----------------------------------------------------------
GBF91741.1_hypothetical_protein_Rsub_04045_[Raphidocelis_subcapitata]                  MPD---------------------------------------------------------
PRW56209.1_hypothetical_protein_C2E21_5275_[Chlorella_sorokiniana]                     M-----------------------------------------------------------
PSC69005.1_hypothetical_protein_C2E20_7401_[Micractinium_conductrix]                   MP----------------------------------------------------------
XP_024543966.1_uncharacterized_protein_LOC112351060_[Selaginella_moellendorffii]       MGLLQL------------------------------------------------------
XP_024376136.1_uncharacterized_protein_LOC112282585_isoform_X3_[Physcomitrella_patens] MPPRQSRNSIAGAFNPMCTPFAAATLACEPENFSMAGTLCPSLSPTL-------------
PNR46676.1_hypothetical_protein_PHYPA_013796_[Physcomitrella_patens]                   MVSC---------CDPLLKGLSMRSTGWESKQ--LASTLIPSLTP---------------
gnl|onekp|CMEQ_scaffold_2011385_Orthotrichum_lyellii                                   ------------------------------------------------------------
gnl|onekp|ORKS_scaffold_2008275_Philonotis_fontana                                     ------------------------------------------------------------
gnl|onekp|HRWG_scaffold_2010347_Buxbaumia_aphylla                                      ------------------------------------------------------------
GAQ79860.1_hypothetical_protein_KFL_000400030_[Klebsormidium_nitens]                   MQAC--------------------------------------------------------
GBG73810.1_hypothetical_protein_CBR_g17148_[Chara_braunii]                             MSSCQLASA-------FCATLDHSLVATSATNNGHVGPTSTSSPPACGLAAVPSVGQSST
XP_006836536.1_uncharacterized_protein_LOC18427423_[Amborella_trichopoda]              MDTVSHT------------------------------------RPSLNPVF---------
NP_194884.1_high_chlorophyll_fluorescence_153_[Arabidopsis_thaliana]                   MARLFVS-----------------------------------------------------
XP_016724527.1_PREDICTED:_uncharacterized_protein_LOC107936335_[Gossypium_hirsutum]    MGSHSII------------------------------TSISSAPPSLPLRFVPEV-----
KAE8038316.1_hypothetical_protein_FH972_010840_[Carpinus_fangiana]                     MASLII------------------------------------------------------
XP_027918900.1_uncharacterized_protein_LOC114177659_[Vigna_unguiculata]                MATFFSV-----------------------------------------------------
XP_019432611.1_PREDICTED:_uncharacterized_protein_LOC109339604_[Lupinus_angustifolius] MATFSLI-----------------------------------------------------
ABK22505.1_unknown_[Picea_sitchensis]                                                  MIDSADIGG------PVSAAMH---------------PSICSHHRSAPLHLIKHV-----
PTQ38080.1_hypothetical_protein_MARPO_0053s0026_[Marchantia_polymorpha]                MATMASTAS---------------------------------------------------

KXZ48069.1_hypothetical_protein_GPECTOR_30g164_[Gonium_pectorale]                      ------------------------------------------------------------
ADL27744.1_cytochrome_c_synthesis_4_protein_[Chlamydomonas_reinhardtii]                ------------------------------------------------------------
GBF91741.1_hypothetical_protein_Rsub_04045_[Raphidocelis_subcapitata]                  ------------------------------------------------------------
PRW56209.1_hypothetical_protein_C2E21_5275_[Chlorella_sorokiniana]                     ------------------------------------------------------------
PSC69005.1_hypothetical_protein_C2E20_7401_[Micractinium_conductrix]                   ------------------------------------------------------------
XP_024543966.1_uncharacterized_protein_LOC112351060_[Selaginella_moellendorffii]       -----------------------------------------------------PSSFSYL
XP_024376136.1_uncharacterized_protein_LOC112282585_isoform_X3_[Physcomitrella_patens] ----------------------------------------LQIECLDFRLSGLSSSS---
PNR46676.1_hypothetical_protein_PHYPA_013796_[Physcomitrella_patens]                   ---------------------------------------------------GFSSSSLRG
gnl|onekp|CMEQ_scaffold_2011385_Orthotrichum_lyellii                                   ------------------------------------------------------------
gnl|onekp|ORKS_scaffold_2008275_Philonotis_fontana                                     ------------------------------------------------------------
gnl|onekp|HRWG_scaffold_2010347_Buxbaumia_aphylla                                      ------------------------------------------------------------
GAQ79860.1_hypothetical_protein_KFL_000400030_[Klebsormidium_nitens]                   ------------------------------------------------------------
GBG73810.1_hypothetical_protein_CBR_g17148_[Chara_braunii]                             LRERSSGRSAWQGPVSTSAVGASSALGASSALLSGDEGGAKRLRRWGDLLGSSELSFAST
XP_006836536.1_uncharacterized_protein_LOC18427423_[Amborella_trichopoda]              -----------------------------------------------------CCNNILF
NP_194884.1_high_chlorophyll_fluorescence_153_[Arabidopsis_thaliana]                   -----------------------------------------------------IPMQPTQ
XP_016724527.1_PREDICTED:_uncharacterized_protein_LOC107936335_[Gossypium_hirsutum]    --------VAFSGHL--------------------------------------SHRKTRG
KAE8038316.1_hypothetical_protein_FH972_010840_[Carpinus_fangiana]                     -----------------------------------------------------SSSVSLT
XP_027918900.1_uncharacterized_protein_LOC114177659_[Vigna_unguiculata]                -----------------------------------------------------SHPISQP
XP_019432611.1_PREDICTED:_uncharacterized_protein_LOC109339604_[Lupinus_angustifolius] -----------------------------------------------------SNSLSNS
ABK22505.1_unknown_[Picea_sitchensis]                                                  ----------W--------------------------------------LPTRMTMKRER
PTQ38080.1_hypothetical_protein_MARPO_0053s0026_[Marchantia_polymorpha]                ----------------------------------------QQLRAFGCAL--------VQ

KXZ48069.1_hypothetical_protein_GPECTOR_30g164_[Gonium_pectorale]                      ------------------------------------------------------------
ADL27744.1_cytochrome_c_synthesis_4_protein_[Chlamydomonas_reinhardtii]                ------------------------------------------------------------
GBF91741.1_hypothetical_protein_Rsub_04045_[Raphidocelis_subcapitata]                  ------------------------------------------------------------
PRW56209.1_hypothetical_protein_C2E21_5275_[Chlorella_sorokiniana]                     ------------------------------------------------------------
PSC69005.1_hypothetical_protein_C2E20_7401_[Micractinium_conductrix]                   ------------------------------------------------------------
XP_024543966.1_uncharacterized_protein_LOC112351060_[Selaginella_moellendorffii]       LYNP------HHRSSAR------------------------SPAFPSGITCSAARVE---
XP_024376136.1_uncharacterized_protein_LOC112282585_isoform_X3_[Physcomitrella_patens] ----------------------AGRGRWA-----VGT----GTCITRQGLAAGAWSD---
PNR46676.1_hypothetical_protein_PHYPA_013796_[Physcomitrella_patens]                   VSNR-----VTMAMEMRRIFARSLRGASH-----VVVAAMATSKPAENGFTADIWSG---
gnl|onekp|CMEQ_scaffold_2011385_Orthotrichum_lyellii                                   -----------------------------------------GPCITRQGFVADVWRG---
gnl|onekp|ORKS_scaffold_2008275_Philonotis_fontana                                     ------------------------------------------------GFAADVWKG---
gnl|onekp|HRWG_scaffold_2010347_Buxbaumia_aphylla                                      --------------------------------------------LLRIGSHENVWSG---
GAQ79860.1_hypothetical_protein_KFL_000400030_[Klebsormidium_nitens]                   -----------------------------------------TSSLRENGRLSSNWSS---
GBG73810.1_hypothetical_protein_CBR_g17148_[Chara_braunii]                             MRRPRR----HQSTGVRVSVCRSRHGDGNHHSGNGGVDKPVRRSFAEISANKPLLLQ---
XP_006836536.1_uncharacterized_protein_LOC18427423_[Amborella_trichopoda]              RHSP---------------------------------YFPPQRRYVEGIGCSPWQLS---
NP_194884.1_high_chlorophyll_fluorescence_153_[Arabidopsis_thaliana]                   ISFP-------------------------------------ASSSQPLLSPPANNFTDGG
XP_016724527.1_PREDICTED:_uncharacterized_protein_LOC107936335_[Gossypium_hirsutum]    LLLY---------------------------------PQTANAFSSPLLVHAPVVFA---
KAE8038316.1_hypothetical_protein_FH972_010840_[Carpinus_fangiana]                     RSQP---------------------------------SIPTRAFFKPPHVSKPPAIR---
XP_027918900.1_uncharacterized_protein_LOC114177659_[Vigna_unguiculata]                SAFA-------------------------------------SFRAAPFFNSPPLRIH---
XP_019432611.1_PREDICTED:_uncharacterized_protein_LOC109339604_[Lupinus_angustifolius] LTFL---------------------------------SPPPSSRAPPFPISPELHFS---
ABK22505.1_unknown_[Picea_sitchensis]                                                  VQFPSL---------------STNNTDILFHSCCGVSRGLPHEGFRFGARRPPVQIA---
PTQ38080.1_hypothetical_protein_MARPO_0053s0026_[Marchantia_polymorpha]                LRRPRVACNVVKIVCFQISDCDLKMAGF-------------SPSSGAFGTKAAARRP---

KXZ48069.1_hypothetical_protein_GPECTOR_30g164_[Gonium_pectorale]                      ------------------TGIEDTIWVNIGSAAALVGATVGATFVGALAIARGIDNA-ED
ADL27744.1_cytochrome_c_synthesis_4_protein_[Chlamydomonas_reinhardtii]                ------------------TGIEDTIWVNVGSAAALVGATIGATFVGAMAISKGIE---AE
GBF91741.1_hypothetical_protein_Rsub_04045_[Raphidocelis_subcapitata]                  --------------PATAAAESSTIWVTAASSVALVVGSMGATVIGAFIASRALDDLDPQ
PRW56209.1_hypothetical_protein_C2E21_5275_[Chlorella_sorokiniana]                     --------------GLPGVAYDGSMWVTAASSVGLVVGSMGLVAGSAALLARRMA---AG
PSC69005.1_hypothetical_protein_C2E20_7401_[Micractinium_conductrix]                   --------------GLAGTAYDGSVWVTAVSSVGLVVASAGFVAAGAALVARRVA---SG
XP_024543966.1_uncharacterized_protein_LOC112351060_[Selaginella_moellendorffii]       RAAAS--RRRRL--VEIGAVSDEKFRTIAAAAVALPIGMAAMAVFGAFLLQRQIR---ME
XP_024376136.1_uncharacterized_protein_LOC112282585_isoform_X3_[Physcomitrella_patens] RIRQM--KSRRVA-DPIRAMGEDTMWITASSAVGLVLSLAAVAALGGYLLSMGVD---NA
PNR46676.1_hypothetical_protein_PHYPA_013796_[Physcomitrella_patens]                   RIRRG--KENKQGVGPIRAMGDETMWITASSAVALVVALAAFAALGGYLLSMGVD---NE
gnl|onekp|CMEQ_scaffold_2011385_Orthotrichum_lyellii                                   RIGRR--RSRKVA-GPIRAMEEETMWITASSAVALVIMLAAFAALGGYLLSVSVD---NE
gnl|onekp|ORKS_scaffold_2008275_Philonotis_fontana                                     RIGRR--RSRKVAPGPVRAMGEETMWITASSAVALVVALAAFAALGGYLLSMGVD---NE
gnl|onekp|HRWG_scaffold_2010347_Buxbaumia_aphylla                                      KGLSL--KIRKSV-GPIRAMGEETMWVTASSAVAVVVVLAACAALGGFLLSMGAE---NE
GAQ79860.1_hypothetical_protein_KFL_000400030_[Klebsormidium_nitens]                   -------LHQVAPLYEIAAGGE---TINPWSLIALAVGTAALAYLSGLVMGKQLE---DE
GBG73810.1_hypothetical_protein_CBR_g17148_[Chara_braunii]                             AGQSR--RRGPASLTTTRALTEDTFWISALQATVVAVGTGAMAALSAFALREGLK---RE
XP_006836536.1_uncharacterized_protein_LOC18427423_[Amborella_trichopoda]              ATSRRRVETKKGVLVVPRAGGGPPSSTILIFAFVFPLSLIAITVLTSIRIADKLD---QQ
NP_194884.1_high_chlorophyll_fluorescence_153_[Arabidopsis_thaliana]                   AGGLCLTRRIRDSSVVTRAG---PSTSSYLLAFAIPATLIAATVFTSIKIADKLD---ED
XP_016724527.1_PREDICTED:_uncharacterized_protein_LOC107936335_[Gossypium_hirsutum]    FSKDLFNRKRRGLEVVTRAG---ANTSSYVFAAVFPLSLLAITIFTSIKIADKLD---ED
KAE8038316.1_hypothetical_protein_FH972_010840_[Carpinus_fangiana]                     FLGNTSRRRTRGLTAVTRAG---PSTSYYIFALALPFSLLAVTIFASIRVADKLD---RD
XP_027918900.1_uncharacterized_protein_LOC114177659_[Vigna_unguiculata]                TGG----RKRRGTAVVARAG---PSTKSILFAIALPSSLLAVTIFSALRMGDKLD---QD
XP_019432611.1_PREDICTED:_uncharacterized_protein_LOC109339604_[Lupinus_angustifolius] AADDSHRRKPRGAMVATRAG---PSTSSLVFAFTLPLSLVAVTVFASIRIADKLD---QK
ABK22505.1_unknown_[Picea_sitchensis]                                                  -------NRRRGTRIRAASG---KNLVTVISAIAVPILLVLVTVIVSIVVSEKLD---RE
PTQ38080.1_hypothetical_protein_MARPO_0053s0026_[Marchantia_polymorpha]                QLLSG--QRKRMT-FSTNALPADTYWITVVQALGVSVFMALAAVISGVVINVGLE---VK

KXZ48069.1_hypothetical_protein_GPECTOR_30g164_[Gonium_pectorale]                      LVDPDLR---AQQNGL--------------------------------------------
ADL27744.1_cytochrome_c_synthesis_4_protein_[Chlamydomonas_reinhardtii]                LLDEDAR---AQMNGA--------------------------------------------
GBF91741.1_hypothetical_protein_Rsub_04045_[Raphidocelis_subcapitata]                  LAEDEAA-----------------------GLAGQEEQ----------------------
PRW56209.1_hypothetical_protein_C2E21_5275_[Chlorella_sorokiniana]                     EIRMPDVNAGMGG-G------------------AQPARR--------------------V
PSC69005.1_hypothetical_protein_C2E20_7401_[Micractinium_conductrix]                   AITFGGEEGTVQTRG------------------GQAARR--------------------V
XP_024543966.1_uncharacterized_protein_LOC112351060_[Selaginella_moellendorffii]       ALRDGEEFIKLVEGG----------------MDEETAKN--------------TIL----
XP_024376136.1_uncharacterized_protein_LOC112282585_isoform_X3_[Physcomitrella_patens] ELSEEEKIESLKQQG----------------LYEKPEP----------------------
PNR46676.1_hypothetical_protein_PHYPA_013796_[Physcomitrella_patens]                   ERGEVEKIEKLKEQG----------------VYENPER----------------------
gnl|onekp|CMEQ_scaffold_2011385_Orthotrichum_lyellii                                   QRDEEDKIKRLKEQG----------------LYEKPEP----------------------
gnl|onekp|ORKS_scaffold_2008275_Philonotis_fontana                                     VRGEEEKIERLKEQG----------------LYEKPEP----------------------
gnl|onekp|HRWG_scaffold_2010347_Buxbaumia_aphylla                                      ERYEEEKVESLKAQG----------------GFEEPEK----------------------
GAQ79860.1_hypothetical_protein_KFL_000400030_[Klebsormidium_nitens]                   AEEMETK---LSEQG----------------ENGEPEPS---------------------
GBG73810.1_hypothetical_protein_CBR_g17148_[Chara_braunii]                             LQREAKEARWMQESGMDTRAWRRAQRRRELGPAEEDERWEENGDDVGGYADE-------Y
XP_006836536.1_uncharacterized_protein_LOC18427423_[Amborella_trichopoda]              FLEELAANQAIVEDE----------------------DG--NDDDNAIFVGD--------
NP_194884.1_high_chlorophyll_fluorescence_153_[Arabidopsis_thaliana]                   FLEDIALNQAIKAAE-------------------KGENG--EGDISL-------------
XP_016724527.1_PREDICTED:_uncharacterized_protein_LOC107936335_[Gossypium_hirsutum]    FLEDISINQAVKEAE---------------DEGDDGDDG--GDDDAISL-----------
KAE8038316.1_hypothetical_protein_FH972_010840_[Carpinus_fangiana]                     YFQELEINQAIREAN----------------EDEDEEEE--EEEDKEDI-----------
XP_027918900.1_uncharacterized_protein_LOC114177659_[Vigna_unguiculata]                WREEMAKMEAAKELD----------------EYDDSDDG--SEDDSMETVQE--------
XP_019432611.1_PREDICTED:_uncharacterized_protein_LOC109339604_[Lupinus_angustifolius] FLEEMAMNEAIMEVD---------------EFDKDNDDD--EDDDVETYLQEEPVFPHAL
ABK22505.1_unknown_[Picea_sitchensis]                                                  FEQEIARKRALMKKL---------------GIQKENLDG---------------------
PTQ38080.1_hypothetical_protein_MARPO_0053s0026_[Marchantia_polymorpha]                AVKEFAE---LDKQG----------------LFKEEDYQLVPDEDFGSFENEEALYI---

KXZ48069.1_hypothetical_protein_GPECTOR_30g164_[Gonium_pectorale]                      ------DAQAEAQPAQRTRLRAED-----VLA---AQEPQEQKPAEG-------------
ADL27744.1_cytochrome_c_synthesis_4_protein_[Chlamydomonas_reinhardtii]                ------EAVNSTTPVQRTRLRAED-----VLARQEQQEKQEQASKEQ-------------
GBF91741.1_hypothetical_protein_Rsub_04045_[Raphidocelis_subcapitata]                  ------QQQQQQQQQQPSRRRRMT--LEEAEAEAAAAAAQGKQQAE--------------
PRW56209.1_hypothetical_protein_C2E21_5275_[Chlorella_sorokiniana]                     QPLRKTSSGSEQGISPPPRAESVD------------------------------------
PSC69005.1_hypothetical_protein_C2E20_7401_[Micractinium_conductrix]                   QPLSKSSAQERQGVQPPARADAVD------------------------------------
XP_024543966.1_uncharacterized_protein_LOC112351060_[Selaginella_moellendorffii]       ------SRIEIIGPDLRSEFRRKRPQIQDKIEQEQQQDEQEQEQEQQTEEPELDTKIQTP
XP_024376136.1_uncharacterized_protein_LOC112282585_isoform_X3_[Physcomitrella_patens] ------TQYEEQGVVTRTRKRRSS------------------------------------
PNR46676.1_hypothetical_protein_PHYPA_013796_[Physcomitrella_patens]                   ------TQYEDQGVTTRTRKRRIS------------------------------------
gnl|onekp|CMEQ_scaffold_2011385_Orthotrichum_lyellii                                   ------TQYKDQGVSTRTRKRRIS------------------------------------
gnl|onekp|ORKS_scaffold_2008275_Philonotis_fontana                                     ------TQYEEQGVGTRSRKRRIS------------------------------------
gnl|onekp|HRWG_scaffold_2010347_Buxbaumia_aphylla                                      ------SQYEEQGVGTSTRNRRI-------------------------------------
GAQ79860.1_hypothetical_protein_KFL_000400030_[Klebsormidium_nitens]                   ------------PEPVRKRNRPIR------------------------------------
GBG73810.1_hypothetical_protein_CBR_g17148_[Chara_braunii]                             DEEEEEEEEDDLPAGRRRRNRPVQ------------------------------------
XP_006836536.1_uncharacterized_protein_LOC18427423_[Amborella_trichopoda]              ----------EKPALSRTRNRPKR------------------------------------
NP_194884.1_high_chlorophyll_fluorescence_153_[Arabidopsis_thaliana]                   ------DDVIQEPVLQRTRNRPKR------------------------------------
XP_016724527.1_PREDICTED:_uncharacterized_protein_LOC107936335_[Gossypium_hirsutum]    ------EEIVQEPVLPRTRNRPKR------------------------------------
KAE8038316.1_hypothetical_protein_FH972_010840_[Carpinus_fangiana]                     -----NISFKKEPALQRTRNRPKR------------------------------------
XP_027918900.1_uncharacterized_protein_LOC114177659_[Vigna_unguiculata]                -----EPALQEEPTLPHARNRPKR------------------------------------
XP_019432611.1_PREDICTED:_uncharacterized_protein_LOC109339604_[Lupinus_angustifolius] NRPDVDTSLQEEPAVPRGRNRPKR------------------------------------
ABK22505.1_unknown_[Picea_sitchensis]                                                  ------AKEEKESVLSRPRNRPKR------------------------------------
PTQ38080.1_hypothetical_protein_MARPO_0053s0026_[Marchantia_polymorpha]                ------FTNKDMAGNRSPRNRPVR--------EAEQQELQEVP-----------------

KXZ48069.1_hypothetical_protein_GPECTOR_30g164_[Gonium_pectorale]                      -----AKK
ADL27744.1_cytochrome_c_synthesis_4_protein_[Chlamydomonas_reinhardtii]                -----ATK
GBF91741.1_hypothetical_protein_Rsub_04045_[Raphidocelis_subcapitata]                  -----QKP
PRW56209.1_hypothetical_protein_C2E21_5275_[Chlorella_sorokiniana]                     -----EAQ
PSC69005.1_hypothetical_protein_C2E20_7401_[Micractinium_conductrix]                   -----EVQ
XP_024543966.1_uncharacterized_protein_LOC112351060_[Selaginella_moellendorffii]       VTMMREEN
XP_024376136.1_uncharacterized_protein_LOC112282585_isoform_X3_[Physcomitrella_patens] ------KD
PNR46676.1_hypothetical_protein_PHYPA_013796_[Physcomitrella_patens]                   ------KY
gnl|onekp|CMEQ_scaffold_2011385_Orthotrichum_lyellii                                   -------E
gnl|onekp|ORKS_scaffold_2008275_Philonotis_fontana                                     -------E
gnl|onekp|HRWG_scaffold_2010347_Buxbaumia_aphylla                                      --------
GAQ79860.1_hypothetical_protein_KFL_000400030_[Klebsormidium_nitens]                   -----REP
GBG73810.1_hypothetical_protein_CBR_g17148_[Chara_braunii]                             ------RE
XP_006836536.1_uncharacterized_protein_LOC18427423_[Amborella_trichopoda]              ------EA
NP_194884.1_high_chlorophyll_fluorescence_153_[Arabidopsis_thaliana]                   ------EV
XP_016724527.1_PREDICTED:_uncharacterized_protein_LOC107936335_[Gossypium_hirsutum]    ------EV
KAE8038316.1_hypothetical_protein_FH972_010840_[Carpinus_fangiana]                     ------AA
XP_027918900.1_uncharacterized_protein_LOC114177659_[Vigna_unguiculata]                ------EA
XP_019432611.1_PREDICTED:_uncharacterized_protein_LOC109339604_[Lupinus_angustifolius] ------EA
ABK22505.1_unknown_[Picea_sitchensis]                                                  ------EE
PTQ38080.1_hypothetical_protein_MARPO_0053s0026_[Marchantia_polymorpha]                -----TDQ
```

Dataset S1B. COX16 protein sequences aligned with putative CCS4 homologs from a representative set of the *Viridiplantae* clade. Transmembrane regions are highlighted in grey; similar charged amino acids motifs are colored in red.

```
NP_680684.1_[Arabidopsis_thaliana]_cox16                                               -MTTIETGQK------------------------------TQKSSP----------SGSG
XP_020519494.1_[Amborella_trichopoda]_cox16                                            -M-VMAATNF------------------------------TSSLKK----------TGIS
gnl|onekp|SGTW_scaffold_2031563_Ginkgo_biloba_cox16                                    -L---------------------------------------EGFRK----------VASE
gnl|onekp|MFTM_scaffold_2014754_Pinus_jeffreyi_cox16                                   ---------------------------------------------------------GTE
gnl|onekp|FLTD_scaffold_2017211_Pteris_ensigormis_cox16                                ------------------------------------------------------------
gnl|onekp|HEGQ_scaffold_2075748_Gymnocarpium_dryopteris_cox16                          EM----------------------------------------------------------
XP_024375750.1_[Physcomitrella_patens]_cox16                                           ------------------------------------------------------------
gnl|onekp|TAVP_scaffold_2076630_Calliergon_cordifolium_cox16                           ------------------------------------------------------------
gnl|onekp|YWNF_scaffold_2048961_Hedwigia_ciliata_cox16                                 ------------------------------------------------------------
OAE26400.1_[Marchantia_polymorpha_subsp._ruderalis]_cox16                              ------------------------------------------------------------
gnl|onekp|JPYU_scaffold_2037236_Marchantia_polymorpha_cox16                            NL----------------------------------------------------------
gnl|onekp|WDCW_scaffold_2044360_Mesotaenium_endlicherianum_cox16                       ------------------------------------------------------------
EFJ16133.1_[Selaginella_moellendorffii]_cox16                                          -MST--------------------------------------------------------
GAQ85474.1_[Klebsormidium_nitens]_cox16                                                -MDQ--------------------------------------------------------
XP_011396124.1_[Auxenochlorella_protothecoides]_cox16                                  -M----------------------------------------------------------
KDD76186.1_[Helicosporidium_sp._ATCC_50920]_cox16                                      -M----------------------------------------------------------
PSC73105.1_[Micractinium_conductrix]_cox16                                             -M----------------------------------------------------------
PRW57852.1_[Chlorella_sorokiniana]_cox16                                               ------------------------------------------------------------
XP_005849331.1_[Chlorella_variabilis]_cox16                                            -MAP--------------------------------------------------------
KAA6428408.1_[Trebouxia_sp._A1-2]_cox16                                                -M----------------------------------------------------------
gnl|onekp|JKKI_scaffold_2006811_Lobomonas_rostrata_cox16                               ------------------------------------------------------------
KXZ56362.1_[Gonium_pectorale]_cox16                                                    ------------------------------------------------------------
XP_001703532.1_[Chlamydomonas_reinhardtii]_cox16                                       -M----------------------------------------------------------
gnl|onekp|QRTH_scaffold_2005535_Chloromonas_perforata_cox16                            ------------------------------------------------------------
gnl|onekp|PRIQ_scaffold_2047349_Pleurastrum_insigne_cox16                              -V----------------------------------------------------------
GAX79751.1_[Chlamydomonas_eustigma]_cox16                                              MM----------------------------------------------------------
XP_013900983.1_[Monoraphidium_neglectum]_cox16                                         -M----------------------------------------------------------
GBF97000.1_[Raphidocelis_subcapitata]_cox16                                            -M----------------------------------------------------------
XP_002501108.1_[Micromonas_commoda]_cox16                                              -MSG--------------------------------------------------------
XP_003064351.1_[Micromonas_pusilla_CCMP1545]_cox16                                     -MAA--------------------------------------------------------
XP_007515412.1_[Bathycoccus_prasinos]_cox16                                            -M----------------------------------------------------------
XP_022838154.1_[Ostreococcus_tauri]_cox16                                              -M----------------------------------------------------------
XP_001415767.1_[Ostreococcus_lucimarinus_CCE9901]_cox16                                -M----------------------------------------------------------
QDZ22244.1_[Chloropicon_primus]_cox16                                                  -M----------------------------------------------------------
XP_024376136.1_uncharacterized_protein_LOC112282585_isoform_X3_[Physcomitrella_patens] -MPPRQSRNSIAGAFNPMCTPFAAATLACEPENFSMAGTLCPSLSPTLLQIEC-LDFRLS
PNR46676.1_hypothetical_protein_PHYPA_013796_[Physcomitrella_patens]                   -MVSC---------CDPLLKGLSMRSTGWESKQ--LASTLIPSLTP--------------
gnl|onekp|CMEQ_scaffold_2011385_Orthotrichum_lyellii                                   ------------------------------------------------------------
gnl|onekp|ORKS_scaffold_2008275_Philonotis_fontana                                     ------------------------------------------------------------
gnl|onekp|HRWG_scaffold_2010347_Buxbaumia_aphylla                                      ------------------------------------------------------------
GAQ79860.1_hypothetical_protein_KFL_000400030_[Klebsormidium_nitens]                   -MQAC-------------------------------------------------------
GBG73810.1_hypothetical_protein_CBR_g17148_[Chara_braunii]                             -MSSCQLASA-------FCATLDHSLVATSATNNGHVGPTSTSSPPA-----CGLAAVPS
XP_006836536.1_uncharacterized_protein_LOC18427423_[Amborella_trichopoda]              -MDT------------------------------------------------------VS
NP_194884.1_high_chlorophyll_fluorescence_153_[Arabidopsis_thaliana]                   -MARLFV--------------------------------------------------SIP
XP_016724527.1_PREDICTED:_uncharacterized_protein_LOC107936335_[Gossypium_hirsutum]    -MGSHSI------------------------------------------------ITSIS
KAE8038316.1_hypothetical_protein_FH972_010840_[Carpinus_fangiana]                     -MASLII------------------------------------------------SSSVS
XP_027918900.1_uncharacterized_protein_LOC114177659_[Vigna_unguiculata]                -MATFF---------------------------------------------------SVS
XP_019432611.1_PREDICTED:_uncharacterized_protein_LOC109339604_[Lupinus_angustifolius] -MATFSL-----------------------------------------------ISNSLS
ABK22505.1_unknown_[Picea_sitchensis]                                                  -MIDSAD----------------------------IGGPVSAAMHP----------SICS
PTQ38080.1_hypothetical_protein_MARPO_0053s0026_[Marchantia_polymorpha]                -MATMAS-----------------------------------------------------
KXZ48069.1_hypothetical_protein_GPECTOR_30g164_[Gonium_pectorale]                      -MPT--------------------------------------------------------
ADL27744.1_cytochrome_c_synthesis_4_protein_[Chlamydomonas_reinhardtii]                -MST--------------------------------------------------------
GBF91741.1_hypothetical_protein_Rsub_04045_[Raphidocelis_subcapitata]                  -MPD--------------------------------------------------------
PRW56209.1_hypothetical_protein_C2E21_5275_[Chlorella_sorokiniana]                     -M----------------------------------------------------------
PSC69005.1_hypothetical_protein_C2E20_7401_[Micractinium_conductrix]                   -MP---------------------------------------------------------
XP_024543966.1_uncharacterized_protein_LOC112351060_[Selaginella_moellendorffii]       -MGLLQLPSS-------FSYLL---------------------YNP--------------

NP_680684.1_[Arabidopsis_thaliana]_cox16                                               TTPTGTL-----------------------------------------------------
XP_020519494.1_[Amborella_trichopoda]_cox16                                            VPSTAQI-----------------------------------------------------
gnl|onekp|SGTW_scaffold_2031563_Ginkgo_biloba_cox16                                    MTATSTAI----------------------------------------------------
gnl|onekp|MFTM_scaffold_2014754_Pinus_jeffreyi_cox16                                   RAPTSTAN----------------------------------------------------
gnl|onekp|FLTD_scaffold_2017211_Pteris_ensigormis_cox16                                ------------------------------------------------------------
gnl|onekp|HEGQ_scaffold_2075748_Gymnocarpium_dryopteris_cox16                          ------------------------------------------------------------
XP_024375750.1_[Physcomitrella_patens]_cox16                                           ------------------------------------------------------------
gnl|onekp|TAVP_scaffold_2076630_Calliergon_cordifolium_cox16                           ---SGEV-----------------------------------------------------
gnl|onekp|YWNF_scaffold_2048961_Hedwigia_ciliata_cox16                                 ------------------------------------------------------------
OAE26400.1_[Marchantia_polymorpha_subsp._ruderalis]_cox16                              ------------------------------------------------------------
gnl|onekp|JPYU_scaffold_2037236_Marchantia_polymorpha_cox16                            --CRAES-----------------------------------------------------
gnl|onekp|WDCW_scaffold_2044360_Mesotaenium_endlicherianum_cox16                       ------------------------------------------------------------
EFJ16133.1_[Selaginella_moellendorffii]_cox16                                          ------------------------------------------------------------
GAQ85474.1_[Klebsormidium_nitens]_cox16                                                ------------------------------------------------------------
XP_011396124.1_[Auxenochlorella_protothecoides]_cox16                                  ------------------------------------------------------------
KDD76186.1_[Helicosporidium_sp._ATCC_50920]_cox16                                      ------------------------------------------------------------
PSC73105.1_[Micractinium_conductrix]_cox16                                             ------------------------------------------------------------
PRW57852.1_[Chlorella_sorokiniana]_cox16                                               ------------------------------------------------------------
XP_005849331.1_[Chlorella_variabilis]_cox16                                            ------------------------------------------------------------
KAA6428408.1_[Trebouxia_sp._A1-2]_cox16                                                ------------------------------------------------------------
gnl|onekp|JKKI_scaffold_2006811_Lobomonas_rostrata_cox16                               ------------------------------------------------------------
KXZ56362.1_[Gonium_pectorale]_cox16                                                    ------------------------------------------------------------
XP_001703532.1_[Chlamydomonas_reinhardtii]_cox16                                       ------------------------------------------------------------
gnl|onekp|QRTH_scaffold_2005535_Chloromonas_perforata_cox16                            ------------------------------------------------------------
gnl|onekp|PRIQ_scaffold_2047349_Pleurastrum_insigne_cox16                              ------------------------------------------------------------
GAX79751.1_[Chlamydomonas_eustigma]_cox16                                              ------------------------------------------------------------
XP_013900983.1_[Monoraphidium_neglectum]_cox16                                         ------------------------------------------------------------
GBF97000.1_[Raphidocelis_subcapitata]_cox16                                            ------------------------------------------------------------
XP_002501108.1_[Micromonas_commoda]_cox16                                              ------------------------------------------------------------
XP_003064351.1_[Micromonas_pusilla_CCMP1545]_cox16                                     ------------------------------------------------------------
XP_007515412.1_[Bathycoccus_prasinos]_cox16                                            ------------------------------------------------------------
XP_022838154.1_[Ostreococcus_tauri]_cox16                                              ------------------------------------------------------------
XP_001415767.1_[Ostreococcus_lucimarinus_CCE9901]_cox16                                ------------------------------------------------------------
QDZ22244.1_[Chloropicon_primus]_cox16                                                  ------------------------------------------------------------
XP_024376136.1_uncharacterized_protein_LOC112282585_isoform_X3_[Physcomitrella_patens] GLSSSS--------------------AGRGRWAVGTGTCIT----RQGLAAGAWSD----
PNR46676.1_hypothetical_protein_PHYPA_013796_[Physcomitrella_patens]                   GFSSSSLRGVSNRVTMAMEMRRIFARSLRGASHVVVAAMATSKPAENGFTADIWSG----
gnl|onekp|CMEQ_scaffold_2011385_Orthotrichum_lyellii                                   ------------------------------------GPCIT----RQGFVADVWRG----
gnl|onekp|ORKS_scaffold_2008275_Philonotis_fontana                                     -----------------------------------------------GFAADVWKG----
gnl|onekp|HRWG_scaffold_2010347_Buxbaumia_aphylla                                      ---------------------------------------LL----RIGSHENVWSG----
GAQ79860.1_hypothetical_protein_KFL_000400030_[Klebsormidium_nitens]                   ---TSSLR-------------------------------------ENGRLSSNW------
GBG73810.1_hypothetical_protein_CBR_g17148_[Chara_braunii]                             VGQSSTLR-------------------------------------ERSSGRSAWQGPVST
XP_006836536.1_uncharacterized_protein_LOC18427423_[Amborella_trichopoda]              HTRPSLNPVFCCN-------------------------------------------NILF
NP_194884.1_high_chlorophyll_fluorescence_153_[Arabidopsis_thaliana]                   MQPTQIS-----------------------------------------------------
XP_016724527.1_PREDICTED:_uncharacterized_protein_LOC107936335_[Gossypium_hirsutum]    SAPPSLPLRFVPE-------------------------------------------VVAF
KAE8038316.1_hypothetical_protein_FH972_010840_[Carpinus_fangiana]                     LTRSQPS-----------------------------------------------------
XP_027918900.1_uncharacterized_protein_LOC114177659_[Vigna_unguiculata]                HPISQPSA----------------------------------------------------
XP_019432611.1_PREDICTED:_uncharacterized_protein_LOC109339604_[Lupinus_angustifolius] NSLTFLSP----------------------------------------------------
ABK22505.1_unknown_[Picea_sitchensis]                                                  HHRSAPLHLI----------------------------------------KHVWLP----
PTQ38080.1_hypothetical_protein_MARPO_0053s0026_[Marchantia_polymorpha]                -TASQQLRAFGC------------------------------------------------
KXZ48069.1_hypothetical_protein_GPECTOR_30g164_[Gonium_pectorale]                      ------------------------------------------------------------
ADL27744.1_cytochrome_c_synthesis_4_protein_[Chlamydomonas_reinhardtii]                ------------------------------------------------------------
GBF91741.1_hypothetical_protein_Rsub_04045_[Raphidocelis_subcapitata]                  ------------------------------------------------------------
PRW56209.1_hypothetical_protein_C2E21_5275_[Chlorella_sorokiniana]                     ------------------------------------------------------------
PSC69005.1_hypothetical_protein_C2E20_7401_[Micractinium_conductrix]                   ------------------------------------------------------------
XP_024543966.1_uncharacterized_protein_LOC112351060_[Selaginella_moellendorffii]       HHRSSAR-----------------------------------------------------

NP_680684.1_[Arabidopsis_thaliana]_cox16                                               ------------------------------------------------------------
XP_020519494.1_[Amborella_trichopoda]_cox16                                            ------------------------------------------------------------
gnl|onekp|SGTW_scaffold_2031563_Ginkgo_biloba_cox16                                    ------------------------------------------------------------
gnl|onekp|MFTM_scaffold_2014754_Pinus_jeffreyi_cox16                                   ------------------------------------------------------------
gnl|onekp|FLTD_scaffold_2017211_Pteris_ensigormis_cox16                                ------------------------------------------------------------
gnl|onekp|HEGQ_scaffold_2075748_Gymnocarpium_dryopteris_cox16                          ------------------------------------------------------------
XP_024375750.1_[Physcomitrella_patens]_cox16                                           ------------------------------------------------------------
gnl|onekp|TAVP_scaffold_2076630_Calliergon_cordifolium_cox16                           ------------------------------------------------------------
gnl|onekp|YWNF_scaffold_2048961_Hedwigia_ciliata_cox16                                 ------------------------------------------------------------
OAE26400.1_[Marchantia_polymorpha_subsp._ruderalis]_cox16                              ------------------------------------------------------------
gnl|onekp|JPYU_scaffold_2037236_Marchantia_polymorpha_cox16                            ------------------------------------------------------------
gnl|onekp|WDCW_scaffold_2044360_Mesotaenium_endlicherianum_cox16                       ------------------------------------------------------------
EFJ16133.1_[Selaginella_moellendorffii]_cox16                                          ------------------------------------------------------------
GAQ85474.1_[Klebsormidium_nitens]_cox16                                                ------------------------------------------------------------
XP_011396124.1_[Auxenochlorella_protothecoides]_cox16                                  ------------------------------------------------------------
KDD76186.1_[Helicosporidium_sp._ATCC_50920]_cox16                                      ------------------------------------------------------------
PSC73105.1_[Micractinium_conductrix]_cox16                                             ------------------------------------------------------------
PRW57852.1_[Chlorella_sorokiniana]_cox16                                               ------------------------------------------------------------
XP_005849331.1_[Chlorella_variabilis]_cox16                                            ------------------------------------------------------------
KAA6428408.1_[Trebouxia_sp._A1-2]_cox16                                                ------------------------------------------------------------
gnl|onekp|JKKI_scaffold_2006811_Lobomonas_rostrata_cox16                               ------------------------------------------------------------
KXZ56362.1_[Gonium_pectorale]_cox16                                                    ------------------------------------------------------------
XP_001703532.1_[Chlamydomonas_reinhardtii]_cox16                                       ------------------------------------------------------------
gnl|onekp|QRTH_scaffold_2005535_Chloromonas_perforata_cox16                            ------------------------------------------------------------
gnl|onekp|PRIQ_scaffold_2047349_Pleurastrum_insigne_cox16                              ------------------------------------------------------------
GAX79751.1_[Chlamydomonas_eustigma]_cox16                                              ------------------------------------------------------------
XP_013900983.1_[Monoraphidium_neglectum]_cox16                                         ------------------------------------------------------------
GBF97000.1_[Raphidocelis_subcapitata]_cox16                                            ------------------------------------------------------------
XP_002501108.1_[Micromonas_commoda]_cox16                                              ------------------------------------------------------------
XP_003064351.1_[Micromonas_pusilla_CCMP1545]_cox16                                     ------------------------------------------------------------
XP_007515412.1_[Bathycoccus_prasinos]_cox16                                            ------------------------------------------------------------
XP_022838154.1_[Ostreococcus_tauri]_cox16                                              ------------------------------------------------------------
XP_001415767.1_[Ostreococcus_lucimarinus_CCE9901]_cox16                                ------------------------------------------------------------
QDZ22244.1_[Chloropicon_primus]_cox16                                                  ------------------------------------------------------------
XP_024376136.1_uncharacterized_protein_LOC112282585_isoform_X3_[Physcomitrella_patens] ------------------------------------------------------------
PNR46676.1_hypothetical_protein_PHYPA_013796_[Physcomitrella_patens]                   ------------------------------------------------------------
gnl|onekp|CMEQ_scaffold_2011385_Orthotrichum_lyellii                                   ------------------------------------------------------------
gnl|onekp|ORKS_scaffold_2008275_Philonotis_fontana                                     ------------------------------------------------------------
gnl|onekp|HRWG_scaffold_2010347_Buxbaumia_aphylla                                      ------------------------------------------------------------
GAQ79860.1_hypothetical_protein_KFL_000400030_[Klebsormidium_nitens]                   ------------------------------------------------------------
GBG73810.1_hypothetical_protein_CBR_g17148_[Chara_braunii]                             SAVGASSALGASSALLSGDEGGAKRLRRWGDLLGSSELSFASTMRRPRRHQSTGVRVSVC
XP_006836536.1_uncharacterized_protein_LOC18427423_[Amborella_trichopoda]              RH----------------------------------------------------------
NP_194884.1_high_chlorophyll_fluorescence_153_[Arabidopsis_thaliana]                   --------------------------------------------------------FPAS
XP_016724527.1_PREDICTED:_uncharacterized_protein_LOC107936335_[Gossypium_hirsutum]    SG------------------------------------H--LSHRKTRGL----LLYPQT
KAE8038316.1_hypothetical_protein_FH972_010840_[Carpinus_fangiana]                     --------------------------------------------------------IPTR
XP_027918900.1_uncharacterized_protein_LOC114177659_[Vigna_unguiculata]                --------------------------------------------------------FASF
XP_019432611.1_PREDICTED:_uncharacterized_protein_LOC109339604_[Lupinus_angustifolius] --------------------------------------------------------PPSS
ABK22505.1_unknown_[Picea_sitchensis]                                                  ---------------------------------------TRMTMKRER------VQFPSL
PTQ38080.1_hypothetical_protein_MARPO_0053s0026_[Marchantia_polymorpha]                ---------------------------------------ALVQLRRPRVACN-VVKIVCF
KXZ48069.1_hypothetical_protein_GPECTOR_30g164_[Gonium_pectorale]                      ------------------------------------------------------------
ADL27744.1_cytochrome_c_synthesis_4_protein_[Chlamydomonas_reinhardtii]                ------------------------------------------------------------
GBF91741.1_hypothetical_protein_Rsub_04045_[Raphidocelis_subcapitata]                  ------------------------------------------------------------
PRW56209.1_hypothetical_protein_C2E21_5275_[Chlorella_sorokiniana]                     ------------------------------------------------------------
PSC69005.1_hypothetical_protein_C2E20_7401_[Micractinium_conductrix]                   ------------------------------------------------------------
XP_024543966.1_uncharacterized_protein_LOC112351060_[Selaginella_moellendorffii]       ------------------------------------------------------------

NP_680684.1_[Arabidopsis_thaliana]_cox16                                               ---------------------------------KQSSASFKRW-----------------
XP_020519494.1_[Amborella_trichopoda]_cox16                                            ---------------------------------KQSASSFKRW-----------------
gnl|onekp|SGTW_scaffold_2031563_Ginkgo_biloba_cox16                                    -------------------------------QRAPKAASYKRW-----------------
gnl|onekp|MFTM_scaffold_2014754_Pinus_jeffreyi_cox16                                   -------------------------------GHTSKIANYKRW-----------------
gnl|onekp|FLTD_scaffold_2017211_Pteris_ensigormis_cox16                                --------------------------------SSTIVKAYRRK-----------------
gnl|onekp|HEGQ_scaffold_2075748_Gymnocarpium_dryopteris_cox16                          --------------------------------STSIAKAYRRK-----------------
XP_024375750.1_[Physcomitrella_patens]_cox16                                           -------------------------------------MAVKRW-----------------
gnl|onekp|TAVP_scaffold_2076630_Calliergon_cordifolium_cox16                           ---------------------------------AMAIASIKRW-----------------
gnl|onekp|YWNF_scaffold_2048961_Hedwigia_ciliata_cox16                                 -------------------------------------AKMKRW-----------------
OAE26400.1_[Marchantia_polymorpha_subsp._ruderalis]_cox16                              ---------------------------------MAGASLYKRM-----------------
gnl|onekp|JPYU_scaffold_2037236_Marchantia_polymorpha_cox16                            ---------------------------------MAGASLYKRM-----------------
gnl|onekp|WDCW_scaffold_2044360_Mesotaenium_endlicherianum_cox16                       ---------------------------------------YRRS-----------------
EFJ16133.1_[Selaginella_moellendorffii]_cox16                                          ---------------------------------VRGIQKGRRL-----------------
GAQ85474.1_[Klebsormidium_nitens]_cox16                                                -------------------------------------VRKRRA-----------------
XP_011396124.1_[Auxenochlorella_protothecoides]_cox16                                  ------------------------------------------------------------
KDD76186.1_[Helicosporidium_sp._ATCC_50920]_cox16                                      ------------------------------------------------------------
PSC73105.1_[Micractinium_conductrix]_cox16                                             ------------------------------------------------------------
PRW57852.1_[Chlorella_sorokiniana]_cox16                                               ------------------------------------------------------------
XP_005849331.1_[Chlorella_variabilis]_cox16                                            ------------------------------------------------------------
KAA6428408.1_[Trebouxia_sp._A1-2]_cox16                                                ------------------------------------------------------------
gnl|onekp|JKKI_scaffold_2006811_Lobomonas_rostrata_cox16                               ------------------------------------------------------------
KXZ56362.1_[Gonium_pectorale]_cox16                                                    ------------------------------------------------------------
XP_001703532.1_[Chlamydomonas_reinhardtii]_cox16                                       ------------------------------------------------------------
gnl|onekp|QRTH_scaffold_2005535_Chloromonas_perforata_cox16                            ------------------------------------------------------------
gnl|onekp|PRIQ_scaffold_2047349_Pleurastrum_insigne_cox16                              ------------------------------------------------------------
GAX79751.1_[Chlamydomonas_eustigma]_cox16                                              ------------------------------------------------------------
XP_013900983.1_[Monoraphidium_neglectum]_cox16                                         -------------------------------------APQQR------------------
GBF97000.1_[Raphidocelis_subcapitata]_cox16                                            -------------------------------------GPPQRE-----------------
XP_002501108.1_[Micromonas_commoda]_cox16                                              -------------------------------------PGTTR------------------
XP_003064351.1_[Micromonas_pusilla_CCMP1545]_cox16                                     -------------------------------------GGLHRA-----------------
XP_007515412.1_[Bathycoccus_prasinos]_cox16                                            ------------------------------------------------------------
XP_022838154.1_[Ostreococcus_tauri]_cox16                                              ------------------------------------------------------------
XP_001415767.1_[Ostreococcus_lucimarinus_CCE9901]_cox16                                ------------------------------------------------------------
QDZ22244.1_[Chloropicon_primus]_cox16                                                  ------------------------------------------------------------
XP_024376136.1_uncharacterized_protein_LOC112282585_isoform_X3_[Physcomitrella_patens] ---------------------------------RIRQMKSRRVA------DPIRAMGEDT
PNR46676.1_hypothetical_protein_PHYPA_013796_[Physcomitrella_patens]                   ---------------------------------RIRRGKENKQG-----VGPIRAMGDET
gnl|onekp|CMEQ_scaffold_2011385_Orthotrichum_lyellii                                   ---------------------------------RIGRRRSRKVA------GPIRAMEEET
gnl|onekp|ORKS_scaffold_2008275_Philonotis_fontana                                     ---------------------------------RIGRRRSRKVA-----PGPVRAMGEET
gnl|onekp|HRWG_scaffold_2010347_Buxbaumia_aphylla                                      ---------------------------------KGLSLKIRKSV------GPIRAMGEET
GAQ79860.1_hypothetical_protein_KFL_000400030_[Klebsormidium_nitens]                   -------------------------------------SSLHQV-------APLYEIAAGG
GBG73810.1_hypothetical_protein_CBR_g17148_[Chara_braunii]                             RSRHGDGNHHSGNGGVDKPVRRSFAEISANKPLLLQAGQSRRRG--PASLTTTRALTEDT
XP_006836536.1_uncharacterized_protein_LOC18427423_[Amborella_trichopoda]              -----SPYF---------PPQRRYVEGIGCSPWQLSATSRRRVETKKGVLVVPRAGGGPP
NP_194884.1_high_chlorophyll_fluorescence_153_[Arabidopsis_thaliana]                   SS---QPLL--------SPPANNFTDGGAG-----GLCLTRRI---RDSSVVTRAG---P
XP_016724527.1_PREDICTED:_uncharacterized_protein_LOC107936335_[Gossypium_hirsutum]    ANAFSSPLLVH------APVVFAFSKD----------LFNRKR---RGLEVVTRAG---A
KAE8038316.1_hypothetical_protein_FH972_010840_[Carpinus_fangiana]                     A--FFKPPHVS------KPPAIRFLGN----------TSRRRT---RGLTAVTRAG---P
XP_027918900.1_uncharacterized_protein_LOC114177659_[Vigna_unguiculata]                RA---APFFNS------PPLRIH--------------TGGRKR---RGTAVVARAG---P
XP_019432611.1_PREDICTED:_uncharacterized_protein_LOC109339604_[Lupinus_angustifolius] RA---PPFPIS------PELHFSAADD----------SHRRKP---RGAMVATRAG---P
ABK22505.1_unknown_[Picea_sitchensis]                                                  STNNTDILFHSCCG--VSRGLPHEGFRFGARRPPVQIANRRRG-------TRIRAA-SGK
PTQ38080.1_hypothetical_protein_MARPO_0053s0026_[Marchantia_polymorpha]                QISDCDLKMAG-----FSPSSGAFGTKAAARRPQLLSGQRKRM------TFSTNALPADT
KXZ48069.1_hypothetical_protein_GPECTOR_30g164_[Gonium_pectorale]                      -------------------------------------------------------GIEDT
ADL27744.1_cytochrome_c_synthesis_4_protein_[Chlamydomonas_reinhardtii]                -------------------------------------------------------GIEDT
GBF91741.1_hypothetical_protein_Rsub_04045_[Raphidocelis_subcapitata]                  --------------------------------------------------PATAAAESST
PRW56209.1_hypothetical_protein_C2E21_5275_[Chlorella_sorokiniana]                     --------------------------------------------------GLPGVAYDGS
PSC69005.1_hypothetical_protein_C2E20_7401_[Micractinium_conductrix]                   --------------------------------------------------GLAGTAYDGS
XP_024543966.1_uncharacterized_protein_LOC112351060_[Selaginella_moellendorffii]       -----------------SPAFPSGITCSAARVERAAASRRRRL-------VEIGAVSDEK

NP_680684.1_[Arabidopsis_thaliana]_cox16                                               --GRRHPFVRYGLPMISLTVFGALGLGQLLQ---------------GSKDIAKVKDDQEW
XP_020519494.1_[Amborella_trichopoda]_cox16                                            --GRKYPFVRYGLPLISLTVLGSVGLGHLLQ---------------GSKEVEKVKDDREW
gnl|onekp|SGTW_scaffold_2031563_Ginkgo_biloba_cox16                                    --GRTNPFLRYGLPLISLTIFGAVGLAHLQQ---------------GRKDMEKVKEDREW
gnl|onekp|MFTM_scaffold_2014754_Pinus_jeffreyi_cox16                                   --GRTNPFLRFGLPLISLTIFGAVGLGHLQQ---------------GRKDMEEVREDREW
gnl|onekp|FLTD_scaffold_2017211_Pteris_ensigormis_cox16                                --VRSSPFLRFGLPLVSLTIVGFYGVGHLVQ---------------GRRDVSNVLDEKGW
gnl|onekp|HEGQ_scaffold_2075748_Gymnocarpium_dryopteris_cox16                          --GRSRPFLQFGLPLVTLTVVGALGIGHLHQ---------------GRRDVAAVLDEQGW
XP_024375750.1_[Physcomitrella_patens]_cox16                                           -----NPFLKFGVPLISLTVLGSVALAHLQQ---------------GRKDVLTARDEKEW
gnl|onekp|TAVP_scaffold_2076630_Calliergon_cordifolium_cox16                           -----NPFIKFGVPLISLTVLGSVALAHLQQ---------------GRKDVLNARDEKEW
gnl|onekp|YWNF_scaffold_2048961_Hedwigia_ciliata_cox16                                 -----NPFIKFGVPLIALTVLGSVALAHLQQ---------------GRKDVLTARDEKEW
OAE26400.1_[Marchantia_polymorpha_subsp._ruderalis]_cox16                              --GRMNPFMRYGMPMITLTVLGSLGLSHLQQ---------------GRKDVANARDDRQW
gnl|onekp|JPYU_scaffold_2037236_Marchantia_polymorpha_cox16                            --GRMNPFMRYGMPMITLTVLGSLGLSHLQQ---------------GRKDVANARDDRQW
gnl|onekp|WDCW_scaffold_2044360_Mesotaenium_endlicherianum_cox16                       --SRGGPFVKYGIPLLSLVVLGYASIGHLMQ---------------GRRDISQAKDDADW
EFJ16133.1_[Selaginella_moellendorffii]_cox16                                          --SGSNRFFLFGLPIIAGSVIGYVGIAQIVA---------------GRTEVVHERDERDW
GAQ85474.1_[Klebsormidium_nitens]_cox16                                                --GGGGPFLRFGLPMTVFSVVGFYGLVHLTQ---------------GRREMINAHSPAQL
XP_011396124.1_[Auxenochlorella_protothecoides]_cox16                                  SKNKSFAFAKAGLPFIVLTVGGWVGLSKTIQ---------------GRIDAQEAQTKEL-
KDD76186.1_[Helicosporidium_sp._ATCC_50920]_cox16                                      -----GHLVRGAFPFVALTVAGWLGVSQILQ---------------GRIDSQEAHQSAV-
PSC73105.1_[Micractinium_conductrix]_cox16                                             APRQAHPFVRAGVPLLGLTIGGFAALQFFIQ---------------GRIDVQDAQRKEL-
PRW57852.1_[Chlorella_sorokiniana]_cox16                                               --------------MLGLTLGGFLGLRFFLQ---------------GRLDVQDAQRKEL-
XP_005849331.1_[Chlorella_variabilis]_cox16                                            PRQGVHPFLRAGVPLLGLTIGGFLGLKFFVQ---------------GRLDVQDAQAREL-
KAA6428408.1_[Trebouxia_sp._A1-2]_cox16                                                -TQGARAFGTVGLPLFVLTVSGFYGLSHLVQ---------------GKFDVQAQRQKVV-
gnl|onekp|JKKI_scaffold_2006811_Lobomonas_rostrata_cox16                               ------SFAMFAVPFGVFMIGGWWGLAQIIQ---------------SKRELKGATSGLR-
KXZ56362.1_[Gonium_pectorale]_cox16                                                    ---------------MLFMIGGWWGLAQLVD---------------SKRQLRGASRGLD-
XP_001703532.1_[Chlamydomonas_reinhardtii]_cox16                                       -AKKQASLLQVGVPFMLFMLGGWYALAHLVD---------------SKRQLQNATRGLD-
gnl|onekp|QRTH_scaffold_2005535_Chloromonas_perforata_cox16                            --KNSASLVRHGLPFLAFLIVGWYGLASVVQ---------------SKRELRNASKGVD-
gnl|onekp|PRIQ_scaffold_2047349_Pleurastrum_insigne_cox16                              -SKSTSQFITHGLPFLGFLLVGWYGLASVVQ---------------SKRELRNASKGVD-
GAX79751.1_[Chlamydomonas_eustigma]_cox16                                              -AKTKASIVSQGLPFFAFIVASWYGLSSLVQ---------------NKRDLGSATRGLD-
XP_013900983.1_[Monoraphidium_neglectum]_cox16                                         --WYSGTFARTALPMFGFVLLCWYGLDQLVS---------------SKLKIRQSVRGYD-
GBF97000.1_[Raphidocelis_subcapitata]_cox16                                            -PWYYGRFARTTLPMFGFVVLCWYGLDQLMA---------------SKLKIRQTVRGYD-
XP_002501108.1_[Micromonas_commoda]_cox16                                              --GSSYPFLQRGLPFFAFMIGGSYGISILLQ---------------GRNEVRDAKADVT-
XP_003064351.1_[Micromonas_pusilla_CCMP1545]_cox16                                     --ATSNHWLRSGVPFVTFMLAGSYGISVVLQ---------------GRNDVRDARADVT-
XP_007515412.1_[Bathycoccus_prasinos]_cox16                                            TKRKQLEFLRLGMPFVAFAVLGSFGLSHIIQ---------------GRLDVKDAKQEVN-
XP_022838154.1_[Ostreococcus_tauri]_cox16                                              -TPAESRFVRHGVPFVALVVIGSFGLQKLVA---------------GRLEARDAARTVE-
XP_001415767.1_[Ostreococcus_lucimarinus_CCE9901]_cox16                                -SRGGARFFAYGAPFVALVAVGASGLARVVG---------------GRLEVRDALGEV--
QDZ22244.1_[Chloropicon_primus]_cox16                                                  ----VSKQLKVGVPMIGVVLVGYLSLTSVLQ---------------GRIKDHDSRTLND-
XP_024376136.1_uncharacterized_protein_LOC112282585_isoform_X3_[Physcomitrella_patens] MWITASSAVGLVLSLAAVAALGGYLLSMGVDNAEL-------------------------
PNR46676.1_hypothetical_protein_PHYPA_013796_[Physcomitrella_patens]                   MWITASSAVALVVALAAFAALGGYLLSMGVDNEER-------------------------
gnl|onekp|CMEQ_scaffold_2011385_Orthotrichum_lyellii                                   MWITASSAVALVIMLAAFAALGGYLLSVSVDNEQR-------------------------
gnl|onekp|ORKS_scaffold_2008275_Philonotis_fontana                                     MWITASSAVALVVALAAFAALGGYLLSMGVDNEVR-------------------------
gnl|onekp|HRWG_scaffold_2010347_Buxbaumia_aphylla                                      MWVTASSAVAVVVVLAACAALGGFLLSMGAENEER-------------------------
GAQ79860.1_hypothetical_protein_KFL_000400030_[Klebsormidium_nitens]                   ETINPWSLIALAVGTAALAYLSGLVMGKQLE-----------------------------
GBG73810.1_hypothetical_protein_CBR_g17148_[Chara_braunii]                             FWISALQATVVAVGTGAMAALSAFALREGLK---------------RELQREAKEARWMQ
XP_006836536.1_uncharacterized_protein_LOC18427423_[Amborella_trichopoda]              SSTILIFAFVFPLSLIAITVLTSIRIADKLD---------------QQFLEELAAN----
NP_194884.1_high_chlorophyll_fluorescence_153_[Arabidopsis_thaliana]                   STSSYLLAFAIPATLIAATVFTSIKIADKLD---------------EDFLEDIALN----
XP_016724527.1_PREDICTED:_uncharacterized_protein_LOC107936335_[Gossypium_hirsutum]    NTSSYVFAAVFPLSLLAITIFTSIKIADKLD---------------EDFLEDISIN----
KAE8038316.1_hypothetical_protein_FH972_010840_[Carpinus_fangiana]                     STSYYIFALALPFSLLAVTIFASIRVADKLD---------------RDYFQELEIN----
XP_027918900.1_uncharacterized_protein_LOC114177659_[Vigna_unguiculata]                STKSILFAIALPSSLLAVTIFSALRMGDKLD---------------QDWREEMAKM----
XP_019432611.1_PREDICTED:_uncharacterized_protein_LOC109339604_[Lupinus_angustifolius] STSSLVFAFTLPLSLVAVTVFASIRIADKLD---------------QKFLEEMAMN----
ABK22505.1_unknown_[Picea_sitchensis]                                                  NLVTVISAIAVPILLVLVTVIVSIVVSEKLD---------------REFEQEIARKR---
PTQ38080.1_hypothetical_protein_MARPO_0053s0026_[Marchantia_polymorpha]                YWITVVQALGVSVFMALAAVISGVVINVGLE---------------VKAVKEFAELDKQ-
KXZ48069.1_hypothetical_protein_GPECTOR_30g164_[Gonium_pectorale]                      IWVNIGSAAALVGATVGATFVGALAIARGIDN-------------AEDLVDPDLRAQQN-
ADL27744.1_cytochrome_c_synthesis_4_protein_[Chlamydomonas_reinhardtii]                IWVNVGSAAALVGATIGATFVGAMAISKGIE---------------AELLDEDARAQMN-
GBF91741.1_hypothetical_protein_Rsub_04045_[Raphidocelis_subcapitata]                  IWVTAASSVALVVGSMGATVIGAFIASRALD----------------DLDPQLAEDEAA-
PRW56209.1_hypothetical_protein_C2E21_5275_[Chlorella_sorokiniana]                     MWVTAASSVGLVVGSMGLVAGSAALLARRMA--------------AGEIRMPDVNAGMG-
PSC69005.1_hypothetical_protein_C2E20_7401_[Micractinium_conductrix]                   VWVTAVSSVGLVVASAGFVAAGAALVARRVA--------------SGAITFGGEEGTVQ-
XP_024543966.1_uncharacterized_protein_LOC112351060_[Selaginella_moellendorffii]       FRTIAAAAVALPIGMAAMAVFGAFLLQRQIRMEALRDGEEFIKLVEGGMDEETAKNTIL-

NP_680684.1_[Arabidopsis_thaliana]_cox16                                               EIIETRKALSRTGPVD----AY--------------------------------------
XP_020519494.1_[Amborella_trichopoda]_cox16                                            EIIQTTRALSRTGPLE----GS--------------------------------------
gnl|onekp|SGTW_scaffold_2031563_Ginkgo_biloba_cox16                                    EAIEITKALSREGPFG-NSVKP--------------------------------------
gnl|onekp|MFTM_scaffold_2014754_Pinus_jeffreyi_cox16                                   EAIERTKALSREGPLG-KFVRP--------------------------------------
gnl|onekp|FLTD_scaffold_2017211_Pteris_ensigormis_cox16                                ERIKETEGLTREGPVG-KNLYR--------------------------------------
gnl|onekp|HEGQ_scaffold_2075748_Gymnocarpium_dryopteris_cox16                          ERIKDTEGLTREGSLG-QNMHR--------------------------------------
XP_024375750.1_[Physcomitrella_patens]_cox16                                           AAMAASKALTREGAVA-SAVAE-----------------------------------AVA
gnl|onekp|TAVP_scaffold_2076630_Calliergon_cordifolium_cox16                           ERMAANKALTREGVVA-GAVAD-----------------------------------AVA
gnl|onekp|YWNF_scaffold_2048961_Hedwigia_ciliata_cox16                                 ERMAANKALTREGVIA-GAVAE-----------------------------------AVA
OAE26400.1_[Marchantia_polymorpha_subsp._ruderalis]_cox16                              EAIAQQRSLSREGPLA-DFESK--------------------------------------
gnl|onekp|JPYU_scaffold_2037236_Marchantia_polymorpha_cox16                            EAIAQQRSLSREGPLA-DFESK--------------------------------------
gnl|onekp|WDCW_scaffold_2044360_Mesotaenium_endlicherianum_cox16                       ARSTASNASSAAI----QARVK--------------------------------------
EFJ16133.1_[Selaginella_moellendorffii]_cox16                                          ELLKVTQALSKEGILKN---FK--------------------------------------
GAQ85474.1_[Klebsormidium_nitens]_cox16                                                Q------PLNKEGE-----LKS--------------------------------------
XP_011396124.1_[Auxenochlorella_protothecoides]_cox16                                  ---------NLHAPAA----KQ--------------------------------------
KDD76186.1_[Helicosporidium_sp._ATCC_50920]_cox16                                      ---------SAHAPVS----KQ--------------------------------------
PSC73105.1_[Micractinium_conductrix]_cox16                                             ---------DLRAPVS----KQ--------------------------------------
PRW57852.1_[Chlorella_sorokiniana]_cox16                                               ---------DLRAPVS----KQ--------------------------------------
XP_005849331.1_[Chlorella_variabilis]_cox16                                            ---------DLRAPVS----KQ--------------------------------------
KAA6428408.1_[Trebouxia_sp._A1-2]_cox16                                                ---------DLKVD------PE--------------------------------------
gnl|onekp|JKKI_scaffold_2006811_Lobomonas_rostrata_cox16                               ---------DVEELDPLERMRRRYGLD------------------------------KEY
KXZ56362.1_[Gonium_pectorale]_cox16                                                    ---------MVEEMDPLERMRRRYGLEDGGAGSG----SSGPG---------RRGRGQTP
XP_001703532.1_[Chlamydomonas_reinhardtii]_cox16                                       ---------MVEELDPMERMRRRYGLREGQAGGGGGAVASKRG---------GSGAAAAA
gnl|onekp|QRTH_scaffold_2005535_Chloromonas_perforata_cox16                            ---------AVEEMDPVERMRRRYGIG---------------G---------AAPAVSAP
gnl|onekp|PRIQ_scaffold_2047349_Pleurastrum_insigne_cox16                              ---------SVEEMDPVERMRRRYGIG---------------G---------AAPVAAPS
GAX79751.1_[Chlamydomonas_eustigma]_cox16                                              ---------SVEEMDPVERMRRRYRMSESDSKG---------------------------
XP_013900983.1_[Monoraphidium_neglectum]_cox16                                         ---------KVEEYDPIERLRRLEGARGGGGSGGGGGNSSGHGSNDSGSGP-GKQPGAAA
GBF97000.1_[Raphidocelis_subcapitata]_cox16                                            ---------KVEEYDPAEAMRRLERGGGGGGDGTGGGSGREGGARRAPRAPRGAAEEAAA
XP_002501108.1_[Micromonas_commoda]_cox16                                              ---------DMRAPSKTQNVRR--------------------------------------
XP_003064351.1_[Micromonas_pusilla_CCMP1545]_cox16                                     ---------DMRAPSRTQTARR--------------------------------------
XP_007515412.1_[Bathycoccus_prasinos]_cox16                                            ---------DFRAPVLAQRKRE--------------------------------------
XP_022838154.1_[Ostreococcus_tauri]_cox16                                              ---------DPRAPARAQRRRR--------------------------------------
XP_001415767.1_[Ostreococcus_lucimarinus_CCE9901]_cox16                                ---------DARAPARTQRVRR--------------------------------------
QDZ22244.1_[Chloropicon_primus]_cox16                                                  ---------TPRIPES----NK--------------------------------------
XP_024376136.1_uncharacterized_protein_LOC112282585_isoform_X3_[Physcomitrella_patens] ------------------------------------------------------------
PNR46676.1_hypothetical_protein_PHYPA_013796_[Physcomitrella_patens]                   ------------------------------------------------------------
gnl|onekp|CMEQ_scaffold_2011385_Orthotrichum_lyellii                                   ------------------------------------------------------------
gnl|onekp|ORKS_scaffold_2008275_Philonotis_fontana                                     ------------------------------------------------------------
gnl|onekp|HRWG_scaffold_2010347_Buxbaumia_aphylla                                      ------------------------------------------------------------
GAQ79860.1_hypothetical_protein_KFL_000400030_[Klebsormidium_nitens]                   ------------------------------------------------------------
GBG73810.1_hypothetical_protein_CBR_g17148_[Chara_braunii]                             E-----SGMDTRAWRRAQRRREL----------------------------------GPA
XP_006836536.1_uncharacterized_protein_LOC18427423_[Amborella_trichopoda]              -----------------QAIVE--------------------------------------
NP_194884.1_high_chlorophyll_fluorescence_153_[Arabidopsis_thaliana]                   -----------------QAIKAA-------------------------------------
XP_016724527.1_PREDICTED:_uncharacterized_protein_LOC107936335_[Gossypium_hirsutum]    -----------------QAVKE--------------------------------------
KAE8038316.1_hypothetical_protein_FH972_010840_[Carpinus_fangiana]                     -----------------QAIRE--------------------------------------
XP_027918900.1_uncharacterized_protein_LOC114177659_[Vigna_unguiculata]                -----------------EAAKEL-------------------------------------
XP_019432611.1_PREDICTED:_uncharacterized_protein_LOC109339604_[Lupinus_angustifolius] -----------------EAIMEV-------------------------------------
ABK22505.1_unknown_[Picea_sitchensis]                                                  ------------------------------------------------------------
PTQ38080.1_hypothetical_protein_MARPO_0053s0026_[Marchantia_polymorpha]                -----------------GLFKE--------------------------------------
KXZ48069.1_hypothetical_protein_GPECTOR_30g164_[Gonium_pectorale]                      ---------------GLDAQAEA----------------------------------QPA
ADL27744.1_cytochrome_c_synthesis_4_protein_[Chlamydomonas_reinhardtii]                ---------------GAEAVNST----------------------------------TPV
GBF91741.1_hypothetical_protein_Rsub_04045_[Raphidocelis_subcapitata]                  ------GLAGQEEQQQQQQQQQQ----------------------------------QPS
PRW56209.1_hypothetical_protein_C2E21_5275_[Chlorella_sorokiniana]                     --------------GGAQPARRV----------------------------------QPL
PSC69005.1_hypothetical_protein_C2E20_7401_[Micractinium_conductrix]                   -------------TRGGQAARRV----------------------------------QPL
XP_024543966.1_uncharacterized_protein_LOC112351060_[Selaginella_moellendorffii]       ---------SRIEIIGPDLRSEF-------------------------------------

NP_680684.1_[Arabidopsis_thaliana]_cox16                                               KPKNT------SIEDELKAMQEKVDIN---------------------TYEYKKIPKL--
XP_020519494.1_[Amborella_trichopoda]_cox16                                            KRKQI------SLEEELKALQKSVDIN---------------------NYEYKRIPKP--
gnl|onekp|SGTW_scaffold_2031563_Ginkgo_biloba_cox16                                    K-----------------------------------------------------------
gnl|onekp|MFTM_scaffold_2014754_Pinus_jeffreyi_cox16                                   KSSKT------NLEDELKALQQKIDIN---------------------NFEYKKIPRP--
gnl|onekp|FLTD_scaffold_2017211_Pteris_ensigormis_cox16                                KNKKI------DLEEELKAMQEKMDIN---------------------AFEYKKIPKP--
gnl|onekp|HEGQ_scaffold_2075748_Gymnocarpium_dryopteris_cox16                          KNKKI------NLEEELKVMQEKMDIN---------------------SFEYKKIPRP--
XP_024375750.1_[Physcomitrella_patens]_cox16                                           RKKSL------DLNEELKDVLNKVNID---------------------DFEYVRVPKL--
gnl|onekp|TAVP_scaffold_2076630_Calliergon_cordifolium_cox16                           KKKPL------NLEEEL----QKVRID---------------------DFDYVRVPKP--
gnl|onekp|YWNF_scaffold_2048961_Hedwigia_ciliata_cox16                                 KKKPL------NLEEEL----NKVKID---------------------DFEYVRVPKP--
OAE26400.1_[Marchantia_polymorpha_subsp._ruderalis]_cox16                              KKKHI------DLEEELKAFQLKVDIN---------------------AFEYKAVPKP--
gnl|onekp|JPYU_scaffold_2037236_Marchantia_polymorpha_cox16                            KKKHI------DLEEELKAFQLKVDIN---------------------AFEYKAVPKP--
gnl|onekp|WDCW_scaffold_2044360_Mesotaenium_endlicherianum_cox16                       RESSV------DLEHELEVLQKKVDID---------------------AFEYKPVPKL--
EFJ16133.1_[Selaginella_moellendorffii]_cox16                                          QRKPT------TLEEELKAIQQQVDIN---------------------NFEYKKVPRP--
GAQ85474.1_[Klebsormidium_nitens]_cox16                                                EGKKL------DLNEELRKLNEKVNIN---------------------DYENKPVPKL--
XP_011396124.1_[Auxenochlorella_protothecoides]_cox16                                  RSKKF------DLHEEMDRLRANVDID---------------------HYELKPVPRL--
KDD76186.1_[Helicosporidium_sp._ATCC_50920]_cox16                                      RAKAF------DLASELERVQSSVDID---------------------DYEYKSVPRP--
PSC73105.1_[Micractinium_conductrix]_cox16                                             RAKKF------NLEEELARLKGEVDIN---------------------NYENVPVPKPKQ
PRW57852.1_[Chlorella_sorokiniana]_cox16                                               RAKKF------NLEEELKRLKGEVDLE---------------------NYENIPVPKPK-
XP_005849331.1_[Chlorella_variabilis]_cox16                                            RAKRF------NLEEELARLKGEVDLE---------------------HYENKPVPRPKQ
KAA6428408.1_[Trebouxia_sp._A1-2]_cox16                                                KQKSF------SLEEELQKLRNSVDIN---------------------NYENKPVPRT--
gnl|onekp|JKKI_scaffold_2006811_Lobomonas_rostrata_cox16                               QPPMS------SLEEELEEMKKKVDIY---------------------NFDYKPVPRT--
KXZ56362.1_[Gonium_pectorale]_cox16                                                    QPEIP------SLEQELEDIRRKVDIY---------------------NFDYKPVPRQ--
XP_001703532.1_[Chlamydomonas_reinhardtii]_cox16                                       ASDVP------SLEAELEAMKGKLDIK---------------------SWDYVPVPRT--
gnl|onekp|QRTH_scaffold_2005535_Chloromonas_perforata_cox16                            KPRIP------TLEEELEDTLSKVNIK---------------------DFDYKPVPRP--
gnl|onekp|PRIQ_scaffold_2047349_Pleurastrum_insigne_cox16                              KPRIP------TLEEELEDTLSKVNIK---------------------DFDYKPVPRP--
GAX79751.1_[Chlamydomonas_eustigma]_cox16                                              SVPVS------SLEDELENVHKSINIY---------------------DFDYKPVPRT--
XP_013900983.1_[Monoraphidium_neglectum]_cox16                                         PSALA------SLEEELAALQASLKID---------------------DFDYKPVPRT--
GBF97000.1_[Raphidocelis_subcapitata]_cox16                                            ARARAAVGGAKSLEEELEELTRGIDLR---------------------TFDYKPVPRT--
XP_002501108.1_[Micromonas_commoda]_cox16                                              KNMAF------DIDAERERTIEELGGL----------------E---KDVDMVPVPRPWE
XP_003064351.1_[Micromonas_pusilla_CCMP1545]_cox16                                     RNRAF------DLDDERERTMDALGVR----------------D-GARDVKMVPVPRPWQ
XP_007515412.1_[Bathycoccus_prasinos]_cox16                                            RRKR-------DEKKNEAQLRETVK----------------------EEYELKEVPRP--
XP_022838154.1_[Ostreococcus_tauri]_cox16                                              ARRAM------DVEGEHRRLIEGRESE--------------------ATYEMKPVWRP--
XP_001415767.1_[Ostreococcus_lucimarinus_CCE9901]_cox16                                AMARF------DAEAERRRLIDAREDR--------------------EAYEMRAVWRP--
QDZ22244.1_[Chloropicon_primus]_cox16                                                  KKGKL------DMEEEIKRAENVVADE----------------------YELKSTRKGG-
XP_024376136.1_uncharacterized_protein_LOC112282585_isoform_X3_[Physcomitrella_patens] -----------SEEEKIESLKQQGLYE---KPEP------------TQYEEQGVVTRTRK
PNR46676.1_hypothetical_protein_PHYPA_013796_[Physcomitrella_patens]                   -----------GEVEKIEKLKEQGVYE---NPER------------TQYEDQGVTTRTRK
gnl|onekp|CMEQ_scaffold_2011385_Orthotrichum_lyellii                                   -----------DEEDKIKRLKEQGLYE---KPEP------------TQYKDQGVSTRTRK
gnl|onekp|ORKS_scaffold_2008275_Philonotis_fontana                                     -----------GEEEKIERLKEQGLYE---KPEP------------TQYEEQGVGTRSRK
gnl|onekp|HRWG_scaffold_2010347_Buxbaumia_aphylla                                      -----------YEEEKVESLKAQGGFE---EPEK------------SQYEEQGVGTSTRN
GAQ79860.1_hypothetical_protein_KFL_000400030_[Klebsormidium_nitens]                   --------------DEAEEMETKLSEQ---GENG-------------EPEPSPEPVRKRN
GBG73810.1_hypothetical_protein_CBR_g17148_[Chara_braunii]                             EEDER----WEENGDDVGGYADEYDEE---EEEE-------------EEDDLPAGRRRRN
XP_006836536.1_uncharacterized_protein_LOC18427423_[Amborella_trichopoda]              ----------------DEDGNDDDNAI-------------------FVGDEKPALSRTRN
NP_194884.1_high_chlorophyll_fluorescence_153_[Arabidopsis_thaliana]                   --------------EKGENGEGDISLD--------------------DVIQEPVLQRTRN
XP_016724527.1_PREDICTED:_uncharacterized_protein_LOC107936335_[Gossypium_hirsutum]    AE---------DEGDDGDDGGDDDAISL------------------EEIVQEPVLPRTRN
KAE8038316.1_hypothetical_protein_FH972_010840_[Carpinus_fangiana]                     ANEDE------DEEEEEEEDKEDINIS---------------------FKKEPALQRTRN
XP_027918900.1_uncharacterized_protein_LOC114177659_[Vigna_unguiculata]                -----------DEYDDSDDGSEDDSMET-VQEEP-------------ALQEEPTLPHARN
XP_019432611.1_PREDICTED:_uncharacterized_protein_LOC109339604_[Lupinus_angustifolius] -----------DEFDKDNDDDEDDDVETYLQEEPVFPHALNRPDVDTSLQEEPAVPRGRN
ABK22505.1_unknown_[Picea_sitchensis]                                                  -----------ALMKKLGIQKENLDGA--------------------KEEKESVLSRPRN
PTQ38080.1_hypothetical_protein_MARPO_0053s0026_[Marchantia_polymorpha]                EDYQL------VPDEDFGSFENEEALY------------------IFTNKDMAGNRSPRN
KXZ48069.1_hypothetical_protein_GPECTOR_30g164_[Gonium_pectorale]                      QRTRL------RAEDVLAAQ---------------------------EPQEQKPAEGAKK
ADL27744.1_cytochrome_c_synthesis_4_protein_[Chlamydomonas_reinhardtii]                QRTRL------RAEDVLARQEQQ------------------------EKQEQASKEQATK
GBF91741.1_hypothetical_protein_Rsub_04045_[Raphidocelis_subcapitata]                  RRRRM------TLEEAEAEAAAAAAQG-------------------KQQAEQKP------
PRW56209.1_hypothetical_protein_C2E21_5275_[Chlorella_sorokiniana]                     RKTSS-------------------------------------------GSEQGISPPPRA
PSC69005.1_hypothetical_protein_C2E20_7401_[Micractinium_conductrix]                   SKSSA-------------------------------------------QERQGVQPPARA
XP_024543966.1_uncharacterized_protein_LOC112351060_[Selaginella_moellendorffii]       RRKRP------QIQDKIEQEQQQDEQE---QEQE-------------QQTEEPELDTKIQ

NP_680684.1_[Arabidopsis_thaliana]_cox16                                               -NESKSS---------------
XP_020519494.1_[Amborella_trichopoda]_cox16                                            -NEGKTGA-------------S
gnl|onekp|SGTW_scaffold_2031563_Ginkgo_biloba_cox16                                    ----------------------
gnl|onekp|MFTM_scaffold_2014754_Pinus_jeffreyi_cox16                                   -SE-------------------
gnl|onekp|FLTD_scaffold_2017211_Pteris_ensigormis_cox16                                ----------------------
gnl|onekp|HEGQ_scaffold_2075748_Gymnocarpium_dryopteris_cox16                          ----------------------
XP_024375750.1_[Physcomitrella_patens]_cox16                                           -KEVAE----------------
gnl|onekp|TAVP_scaffold_2076630_Calliergon_cordifolium_cox16                           ----------------------
gnl|onekp|YWNF_scaffold_2048961_Hedwigia_ciliata_cox16                                 -KE-------------------
OAE26400.1_[Marchantia_polymorpha_subsp._ruderalis]_cox16                              -RQAEINK-------------B
gnl|onekp|JPYU_scaffold_2037236_Marchantia_polymorpha_cox16                            -RQ-------------------
gnl|onekp|WDCW_scaffold_2044360_Mesotaenium_endlicherianum_cox16                       -KD-------------------
EFJ16133.1_[Selaginella_moellendorffii]_cox16                                          -GEKADK---------------
GAQ85474.1_[Klebsormidium_nitens]_cox16                                                -SSDE-----------------
XP_011396124.1_[Auxenochlorella_protothecoides]_cox16                                  -PDEDE----------------
KDD76186.1_[Helicosporidium_sp._ATCC_50920]_cox16                                      -PEEE-----------------
PSC73105.1_[Micractinium_conductrix]_cox16                                             LPEDDDA---------------
PRW57852.1_[Chlorella_sorokiniana]_cox16                                               LPDDDDD---------------
XP_005849331.1_[Chlorella_variabilis]_cox16                                            LAEGEDG---------------
KAA6428408.1_[Trebouxia_sp._A1-2]_cox16                                                -EDE------------------
gnl|onekp|JKKI_scaffold_2006811_Lobomonas_rostrata_cox16                               -EDD------------------
KXZ56362.1_[Gonium_pectorale]_cox16                                                    -AEDDE----------------
XP_001703532.1_[Chlamydomonas_reinhardtii]_cox16                                       -DEDE-----------------
gnl|onekp|QRTH_scaffold_2005535_Chloromonas_perforata_cox16                            -EEDE-----------------
gnl|onekp|PRIQ_scaffold_2047349_Pleurastrum_insigne_cox16                              -EEDE-----------------
GAX79751.1_[Chlamydomonas_eustigma]_cox16                                              -DEEEE----------------
XP_013900983.1_[Monoraphidium_neglectum]_cox16                                         -VDDEE----------------
GBF97000.1_[Raphidocelis_subcapitata]_cox16                                            -EDDG-----------------
XP_002501108.1_[Micromonas_commoda]_cox16                                              EPGAKRRRWW-KGVKRWWG-GA
XP_003064351.1_[Micromonas_pusilla_CCMP1545]_cox16                                     EPGANGRRWW-GGVRRWLKRGE
XP_007515412.1_[Bathycoccus_prasinos]_cox16                                            -TGS----WW------------
XP_022838154.1_[Ostreococcus_tauri]_cox16                                              -PGVEGGH--------------
XP_001415767.1_[Ostreococcus_lucimarinus_CCE9901]_cox16                                -EGGQGH---------------
QDZ22244.1_[Chloropicon_primus]_cox16                                                  ----------------------
XP_024376136.1_uncharacterized_protein_LOC112282585_isoform_X3_[Physcomitrella_patens] RRSSKD----------------
PNR46676.1_hypothetical_protein_PHYPA_013796_[Physcomitrella_patens]                   RRISKY----------------
gnl|onekp|CMEQ_scaffold_2011385_Orthotrichum_lyellii                                   RRISE-----------------
gnl|onekp|ORKS_scaffold_2008275_Philonotis_fontana                                     RRISE-----------------
gnl|onekp|HRWG_scaffold_2010347_Buxbaumia_aphylla                                      RRI-------------------
GAQ79860.1_hypothetical_protein_KFL_000400030_[Klebsormidium_nitens]                   RPIRREP---------------
GBG73810.1_hypothetical_protein_CBR_g17148_[Chara_braunii]                             RPVQRE----------------
XP_006836536.1_uncharacterized_protein_LOC18427423_[Amborella_trichopoda]              RPKREA----------------
NP_194884.1_high_chlorophyll_fluorescence_153_[Arabidopsis_thaliana]                   RPKREV----------------
XP_016724527.1_PREDICTED:_uncharacterized_protein_LOC107936335_[Gossypium_hirsutum]    RPKREV----------------
KAE8038316.1_hypothetical_protein_FH972_010840_[Carpinus_fangiana]                     RPKRAA----------------
XP_027918900.1_uncharacterized_protein_LOC114177659_[Vigna_unguiculata]                RPKREA----------------
XP_019432611.1_PREDICTED:_uncharacterized_protein_LOC109339604_[Lupinus_angustifolius] RPKREA----------------
ABK22505.1_unknown_[Picea_sitchensis]                                                  RPKREE----------------
PTQ38080.1_hypothetical_protein_MARPO_0053s0026_[Marchantia_polymorpha]                RPVREAEQQELQEVPT----DQ
KXZ48069.1_hypothetical_protein_GPECTOR_30g164_[Gonium_pectorale]                      ----------------------
ADL27744.1_cytochrome_c_synthesis_4_protein_[Chlamydomonas_reinhardtii]                ----------------------
GBF91741.1_hypothetical_protein_Rsub_04045_[Raphidocelis_subcapitata]                  ----------------------
PRW56209.1_hypothetical_protein_C2E21_5275_[Chlorella_sorokiniana]                     ESVDEAQ---------------
PSC69005.1_hypothetical_protein_C2E20_7401_[Micractinium_conductrix]                   DAVDEVQ---------------
XP_024543966.1_uncharacterized_protein_LOC112351060_[Selaginella_moellendorffii]       TPVTMMREEN------------
```
