## Supplemental Material for "TWO DISULFIDE-REDUCING PATHWAYS ARE REQUIRED FOR THE MATURATION OF PLASTID *C*-TYPE CYTOCHROMES IN *CHLAMYDOMONAS REINHARDTII*"

**Ankita Das^*†^, Nitya Subrahmanian^*,1^, Stéphane T. Gabilly**^*,2^**, Ekaterina P. Andrianova**^§^**, Igor B. Zhulin**^§^**, Ken Motohashi^‡^, Patrice Paul Hamel^**^.**

*The Ohio State University, Department of Molecular Genetics, 500 Aronoff Laboratory, 318 W. 12^th^ Avenue, Columbus, OH, 43210, USA.

**^†^** Molecular Genetics Graduate Program, The Ohio State University, Columbus, Ohio, USA.

**^‡^** Kyoto Sangyo University, Department of Frontier Life Sciences, Karigamo Motoyama, Kita-ku, Kyoto 603-8047, Japan

^§^ The Ohio State University, Department of Microbiology and Translational Data Analytics Institute 318 W. 12^th^ Avenue, Columbus, OH, 43210, USA.

**^**^** The Ohio State University, Department of Molecular Genetics and Department of Biological Chemistry and Pharmacology, 500 Aronoff Laboratory, 318 W. 12^th^ Avenue, Columbus, OH, 43210, USA.

**Current address**

^1^University of Florida, Department of Neurology, 1275 Center drive, Gainesville, Florida, USA

^2^ Calixar, Bâtiment Laennec, 60 avenue Rockfeller, Lyon, France

**Correspondence**

Patrice P. Hamel

The Ohio State University, Department of Molecular Genetics and Department of Biological Chemistry and Pharmacology.

500 Aronoff Laboratory, 318 W. 12^th^ Avenue, Columbus, OH, 43210, USA.

**SUPPLEMENTAL MATERIAL**

**Supplemental Figures**

Figure S1. Suppressors of Δccs4ccs5 are restored for cytochrome f assembly.

Figure S2. Gain-of-function mutations in CCS4 are dominant.

Figure S3. Conservation of CCS4 in chlorophytes.

**Supplemental Table**

Table S1. List of algal strains used in this study

**Supplemental data**

Data S1. Search for CCS4 ortholog using PSI-BLAST: Conservation of CCS4 in the green lineage and evolutionary relationship to COX16.

Dataset 1A: Multiple Sequence Alignment: CCS4 homologs from a representative set of the Viridiplantae clade,

Dataset 1B: Multiple Sequence Alignment: COX16 protein sequences aligned with putative CCS4 homologs from a representative set of the Viridiplantae clade.

**Figure S1. Suppressors of Δccs4ccs5 are restored for cytochrome f assembly.** Immunoblotting was performed on membrane fractions of total protein extracts prepared from cells grown mixotrophically (on acetate with 0.3 µmol/m^2^/s of light) at 25°C for 5 -7 days. Strains are as in Figure 4C. Samples (100%) correspond to 10 µg of chlorophyll were used and immunodetection with antisera against cyt f. α-PsbO was used as loading control.

**Figure S2. Gain-of-function mutations in CCS4 are dominant.** Ten-fold dilution series of diploids (+/+, ccs5 /ccs5, ccs5/ccs5 CCS4-2 and ccs5/ccs5 CCS4-3) were plated on minimal medium and grown phototrophically (with 30-50 µmol/m^2^/sec of light) at 25°C for 14-18 days (left panel) and mixotrophically in 0.3 µmol/m^2^/sec for 20-21 days (right panel). Two representatives of each diploid constructed with a ccs5 strain are shown.

**Figure S3. Conservation of CCS4 in chlorophytes.** Alignment of CCS4 from Chlamydomonas reinhardtii (accession no. ADL27744.1), Gonium pectoral (accession no. KXZ48069.1), Volvox carteri (accession no. FD920844.1), Dunaliella salina (accession no. BM447122.1) was performed using Clustal omega (Sievers and Higgins 2014). The putative transmembrane domain is highlighted in purple (using TMHMM-2.0 prediction tool) (Krogh et al. 2001). Strictly conserved or similar amino acids are highlighted in yellow. The percentage of similarity to the Chlamydomonas ortholog is indicated. The percentage of charged residues (lysine, arginine, aspartate, and glutamate) in the stroma facing domain is provided.

**Table S1. List of Chlamydomonas strains used in this study**

| **Strains (aliases)** | **Genotype** | **Chlamydomonas Collection center** | **Reference** |
| --- | --- | --- | --- |
| 137C | mt^-^; nit1; nit2 | CC-124 | (Pröschold et al. 2005) |
| 4C^-^ | mt^-^; arg7-8 | CC-5590 | (Subrahmanian et al. 2020) |
| CMJ030 | mt^-^; cw_15_ | CC-4533 | (Li et al. 2019) |
| T78.15b- | mt^-^; ccs5::ARG7Φ | CC-4129 | (Gabilly et al. 2010) |
| ccs5(13) | mt^+^; ccs5::ARG7Φ; cw_15_ | CC-5922 | This work |
| ccs5-2 | mt^-^; cw_15_; ccs5::APHVIII | CC-0000 | This work |
| PH39-35 | mt^-^; ccs5::APHVII; arg7 | CC-5923 | This work |
| PH39-41 | mt^+^; ccs5::APHVIII; arg7 | CC-5924 | This work |
| ccs5(CCS5) | mt^-^; ccs5::ARG7Φ; APHVII; CCS5 | CC-4527 | (Gabilly et al. 2010) |
| ccs4.1 | mt^-^; ccs4-F2D8; arg7-8 | CC-4519 | (Gabilly et al. 2011) |
| ccs4(pCB412) | mt^-^; ccs4-F2D8; ARG7 | - | (Gabilly et al. 2011) |
| ccs4cw_15_ | mt^-^; ccs4-F2D8; cw_15_; arg7-8 | CC-4525 | (Gabilly et al. 2011) |
| ccs4(28) | mt^+^; ccs4-F2D8; cw_15_ | CC-5925 | This work |
| ccs4(pSL18) | mt^-^; ccs4-F2D8; arg7-8; APHVIII | CC-4520 | (Gabilly et al. 2011) |
| ccs4(CCS4) | mt^-^; ccs4-F2D8; arg7-8; APHVIII; CCS4 | CC-4522 | (Gabilly et al. 2011) |
| ccs4(CCDA) | mt^-^; ccs4-F2D8; cw_15_; arg7-8; APHVIII; CCDA | CC-5926 | This work |
| ccs4ccs5(4) | mt^-^; ccs5::ARG7Φ; ccs4-F2D8; APHVIII | CC-4517 | This work |
| ccs4ccs5(5) | mt^-^; ccs5::ARG7Φ; ccs4-F2D8; APHVIII | CC-4518 | This work |
| ccs4ccs5(35) | mt^-^; ccs5::ARG7Φ; ccs4-F2D8; cw_15_ | CC-5927 | This work |
| ccs4ccs5(9) | mt^+^; ccs5::ARG7Φ; ccs4-F2D8; APHVIII | see M&M | This work |
| ccs4ccs5(11) | mt^+^; ccs5::ARG7Φ; ccs4-F2D8; APHVIII | see M&M | This work |
| SU9 | mt^+^; ccs5::ARG7Φ; CCS4-2 | CC-5928 | This work |
| SU11 | mt^+^; ccs5::ARG7Φ; CCS4-3 | CC-5929 | This work |
| PH24-18 | mt^-^; ccs5::ARG7Φ; CCS4-3 | CC-5930 | This work |
| PH46-1M | mt^+^; ccs5::APHVIII; CCS4-2 arg7-8 | CC-5931 | This work |
| CCS4-WT | mt^+^; ccs5::ARG7Φ; ccs4-F2D8; APHVII; CCS4 | CC-5932 | This work |
| CCS4-SU9 | mt^+^; ccs5::ARG7Φ; ccs4-F2D8;APHVII;CCS4-2 | CC-5933 | This work |
| CCS4-SU11 | mt^+^; ccs5::ARG7Φ; ccs4-F2D8;APHVII;CCS4-3 | CC-5934 | This work |
| ΔpetA | mt^+^; petA::aadA | CC-5935 | (Zhou et al. 1996) |
| ccs5 / ccs5 | mt^-^; ccs5::ARG7Φ / mt^+^; ccs5::APHVIII; arg7-8 | - | This work |
| ccs5 / ccs5 CCS4-2 | mt^-^; ccs5::ARG7Φ / mt^+^; ccs5::APHVIII; CCS4-2; arg7-8 | - | This work |
| ccs5 / ccs5 CCS4-2 | mt^+^; ccs5::APHVIII; arg7-8 / mt^-^; ccs5::ARG7Φ; CCS4-3 | - | This work |

**Data S1. Search for CCS4 ortholog using PSI-BLAST**: Using the full protein sequence of cytochrome c synthesis 4 (CCS4) protein from Chlamydomonas reinhardtii (accession ADL27744.1) as a query, we carried out a similarity search using the Basic Alignment Search Tool (BLAST) (Altschul et al. 1997) against the NCBI non-redundant database and detected only two similar sequences in closely related Chlamydomonas eustigma (GAX77750.1) and Gonium pectorale (KXZ48069.1). Searches against Pfam (El-Gebali et al. 2019) and CDD (Lu et al. 2020) databases using HMMER (Potter et al. 2018) and RPS-BLAST, respectively, failed to identify any known domains in these sequences and highly sensitive profile-profile searches using HHpred against all databases implemented in the MPI Bioinformatics ToolKit (Zimmermann et al. 2018) resulted in no significant hits. The search for transmembrane regions and signal peptides revealed that all three proteins were predicted to have a single N-terminal transmembrane helix. We also noticed that all three proteins had a similar length (~ 100 amino acid residues) and an unusually high (>30%) content of charged amino acids in their C-terminal part, which was predicted to reside in a cytoplasm.

To identify distant homologs, we performed exhaustive PSI-BLAST searches (Altschul et al. 1997) initiated with all three sequences against Viridiplantae genomes in the NCBI non-redundant database. Newly found sequences were accessed to match the following criteria: (i) sequence length should not exceed 300 amino acid residues, (ii) contains a single transmembrane helix, and (iii) >30% of the predicted soluble part of the protein is composed of the charged residues. Sequences matching these criteria were used as queries in BLAST and exhaustive PSI-BLAST searches against the following databases: individual Viridiplantae genomes, the entire plant clade in the NCBI non-redundant database, and the 1000 Plant Genome Project (1KP) database (Matasci et al. 2014)

A PSI-BLAST (Position-Specific Iterative Basic Local Alignment Search Tool) initiated with the sequence from Gonium pectorale (KXZ48069.1) retrieved potential homologs in green alga Raphidocelis subcapitata (GBF91741.1) and in moss Physcomitrella patens (accession PNR46676.1 and XP_024376134.1). PSI-BLAST searches initiated with sequences from moss retrieved a final set of hits (with e-value < 8e-09). This set included proteins from C. reinhardtii (ADL27744.1), C. eustigma (GAX77750.1), G. pectorale (KXZ48069.1), R. subcapitata (GBF91741.1), Chara braunii (GBG73810.1), Marchantia polymorpha (PTQ38080.1), P. patens (XP_024376137.1 and PNR46676.1), Selaginella moellendorffii (XP_024543966.1), Picea sitchensis (ABK22505.1), Klebsormidium nitens (GAQ79860.1) and sequences from numerous land plantes, including Arabidopsis thaliana (NP_194884.1). BLAST searches against the 1KP database using sequences from G. pectorale (KXZ48069.1), M. polymorpha (PTQ38080.1) and R. subcapitata GBF91741.1 retrieved two more sequences from mosses, Orthotrichum lyellii (CMEQ_scaffold_2011385) and cf. Physcomicromitrium sp. (YEPO_scaffold_2010104), that were added to the final data set. All sequences from the final set of CCS4 homologs were aligned and the multiple sequence alignment (see Dataset S1A) was used to generate a sequence logo.

BLAST search using G. pectorale (KXZ48069.1) sequence against the genome of Chloromonas perforate retrieved a protein (QRTH_scaffold_2005535), which contained a Pfam domain PF14138 (cytochrome c oxidase assembly protein COX16). In animal cells, COX16 is a small membrane-anchored mitochondrial protein, which participates in the assembly of a cytochrome c oxidase, a respiratory complex but the precise role of COX16 in this process is not well understood (Carlson et al. 2003). Considering the role of CCS4 in the assembly of a photosynthetic complex in C. reinhardtii, we explored this potential remote relationship further. First, we collected COX16 homologs from representative genomes from each major clade of green plants using Chloromonas perforate COX16 as a query in BLAST searches against NCBI non-redundant and 1KP databases. Retrieved COX16 protein sequences were aligned together with putative CCS4 homologs identified in the previous step (Dataset S1B).

All collected protein sequences identified as COX16 homologs were subjected to transmembrane helix searches using Phobius (Käll et al. 2004). Signal peptide or transmembrane region (often overlapping) were predicted in all COX16 sequences. Finally, while analyzing the sequence logo of COX16 Pfam model and our sequence alignments (Figure 6B), we noticed that members of COX16 family had the same key features as CCS4 homologs identified earlier, i.e., the presence of predicted of TM region (or signal peptide in some cases) in the N-terminus and a highly charged C-terminal part and a putative shared motif.

A meaningful phylogenetic analysis of CCS4 proteins cannot be produced due to a short length of the protein sequence and extreme sequence variation. However, several facts suggest that CCS4 and COX16 might be homologs: (i) they share distinct sequence features, such as sequence length, a single transmembrane helix/signal peptide, and a highly charged C-terminus and (ii) they share a common biological function - both are involved in a disulfide reducing pathway. It is plausible that CCS4 might have appeared because of COX16 gene duplication (or they share a common ancestor) in the last universal common ancestor of the Viridiplantae clade. After duplication, one gene copy is usually redundant and thus freed from functional constraints, allowing it to diverge in function or to become a pseudogene (Lynch and Conery 2000). Considerable divergence among CCS4 homologs suggests the possibility of their broader functional diversification in major plant clades (flowering plants, bryophytes, charophytes and chlorophytes) or even at the family level.
